# Structural heterogeneity in mRNA-LNP subpopulations revealed by AF4-SAXS: implications for cargo loading and cell transfection

**DOI:** 10.64898/2026.01.16.699683

**Authors:** Adrian Sanchez-Fernandez, Keira A. Donnelly, Hans Bolinsson, Anna-Maria Börjesdotter, Thomas Rønnemoes Bobak, Simon Erlendsson, Meysam Mohammadi-Zerankeshi, Khaled AboulFotouh, Mohammed R. Kawelah, Fátima Herranz-Trillo, Herje Schagerlöf, Umberto Capasso Palmiero, Kasper Huus, Keith P. Johnston, Alexander E. Marras, Lars Nilsson

## Abstract

Lipid nanoparticles are the leading platform for the delivery of nucleic acid therapeutics, yet their structural complexity remains a significant barrier to achieve rational design and predictable function. Part of this complexity arises from the non-equilibrium assemblies that are difficult to identify using ensemble average techniques given the substantial heterogeneity in all properties. Aiming to overcome the limitations of traditional characterization methods, we combined asymmetric flow field–flow fractionation with in-line small-angle X-ray scattering and spectroscopic analyses, nanoflow cytometry, and cryo-EM to construct detailed structural models of mRNA-loaded nanoparticles formulated with different amounts of mRNA loading (N/P ratios of 3 and 6). This combination of techniques revealed that microfluidic formulation produces structurally diverse nanoparticle subpopulations differing in size, anisotropy, and cargo loading. Notably, these variations extend to the particle internal organization: spheroidal geometries display densely loaded mRNA cores, whereas bleb-like morphologies exhibit reduced mRNA content relative to the lipid amount within segregated domains at the core. NanoFCM further shows that the N/P ratio modulates cargo distribution across individual nanoparticles, with N/P=6 yielding a more uniform mRNA copy number per particle across subpopulations than N/P=3. These differences resulted in higher transfection efficacies for the N/P=6 formulation, highlighting core organization and loading homogeneity as key parameters for efficacious delivery. Together, these results establish a direct link between LNP architecture, internal organization, cargo distribution, and transfection efficiency, underscoring the importance of accounting for heterogeneity in the rational design of nucleic acid delivery systems.

## Introduction

Enhancing cargo delivery efficiency is essential to unlock the full potential of nucleic acid therapeutics, which utilize various RNA classes that can selectively target inherent deficiencies for the treatment or prevention of disease.^1–3^ Recent years have witnessed the emergence of LNPs as the gold-standard vehicles for nucleic acid delivery, enabling the development of the next generation therapies.^4,5^ In this regard, mRNA vaccines have shown exceptional clinical success, most prominently during the COVID-19 global pandemic,^6^ owning to the possibility of rationally engineering these soft nanomaterials to achieve effective payload protection and intracellular delivery. The conventional view describes LNP architecture as a core–shell spheroid, where a dense core contains nucleic acids complexed with ionizable lipids encapsulated within an outer shell of helper and PEG-lipids, and cholesterol distributed across the nanostructure.^7,8^ LNPs are typically formed through rapid microfluidic mixing of lipids in ethanol with acidified aqueous nucleic acid solutions, yielding kinetically arrested assemblies.^7,9,10^ Due to the non-equilibrium nature of this process, their structural and functional properties result from a complex interplay between lipid composition, cargo characteristics, and formulation and storage conditions.^11–13^

While LNP structure and dynamics have been linked to efficacy at multiple stages of delivery—including cellular internalization, endosomal escape, and protein expression—^5,14–16^ our ability to construct accurate structural models remains constrained by the limited resolution and interpretability of existing analytical techniques.^17–19^ Common methods for morphological characterization often lack the sensitivity to distinguish subpopulations or to provide statistically meaningful data into heterogeneous assemblies. For example, ensemble-averaging techniques such as dynamic light scattering (DLS) and small-angle X-ray scattering (SAXS) obscure the contribution of minor yet structurally distinct subpopulations, including “bleb-like” morphologies, which occur at lower abundance than conventional spheroidal particles.^8^ Conversely, single-particle approaches such as cryogenic electron microscopy (cryo-EM) can reveal these unconventional structures but often fail to capture overall population diversity and quantitative distribution. Yet, these unconventional assemblies have been shown to significantly influence delivery efficacy.^8,16^ Unlike the simplified notion of a rather homogeneous population of spherical LNPs, the current picture suggests a much more complex structural landscape, characterized by overall heterogeneity, formation of internal domains, incomplete encapsulation, and local variations in lipid density.^20–24^ In fact, a recent study demonstrated that both the characteristics of lipid constituents and formulation protocol impact the system’s heterogeneity, ultimately affecting delivery efficacy.^25^ Therefore, advancing characterization strategies that combine subpopulation-level resolution with population-scale quantification is essential to achieve a comprehensive understanding of LNP structure–function relationships, ultimately enabling the development of more predictive and rationally designed formulations.^12,13,16,26^

An important parameter in LNP design is the N/P ratio—defined as the molar ratio of protonable amine groups (N) in the ionizable lipid to the phosphate groups (P) in the nucleic acid cargo—which strongly influences particle structure and delivery efficacy.^27^ Systematic studies have shown that varying N/P ratios affects the relative content of lipophilic versus non-lipophilic lipid–RNA complexes during LNP formation, leading to measurable differences in particle loading, population structure, and cytosolic internalization.^28^ In addition, N/P values dictate the degree of internal ordering at the particle core, often yielding more ordered domains at lower ratios and affecting cell transfection efficiency.^29^ This structural transition was further confirmed by computational methods, showing that lower N/P values yield higher internal order and reduced mRNA mobility, whereas at high ratios the core becomes more amorphous.^30^ These observations underscore the structural complexity introduced by variations in N/P ratio, highlighting how these can reshape particle architecture, ultimately complicating efforts to establish general structure–function relationships. However, even formulations with similar external morphology, e.g. comparable size distributions, can harbor distinct internal architectures and delivery efficacies at different N/P ratios, further motivating systematic investigations beyond traditional methods that lack the resolution to account for these changes.

In this study, we challenge the conventional approach to characterize mRNA-loaded LNPs as a single, uniform population and instead reveal a distribution of structural subpopulations that depends sensitively on the N/P ratio. We employed size-based fractionation to resolve structural differences among LNP subpopulations and to elucidate how these relate to internal architecture and cargo distribution. By coupling asymmetric flow field–flow fractionation (AF4) with SAXS,^31,32^ and integrating these data with spectroscopy, cryo-EM, and nano-flow cytometry (NanoFCM) analyses, we show that microfluidic preparation of technologically relevant formulations (N/P ratios 6 and 3) yields heterogeneous systems consisting of two main structural sub-populations: prolate spheroidal assemblies with subtle asymmetry and bleb-like, anisotropic structures. Our results confirm strong correlations between overall morphology, particle density, internal ordering, and cargo loading, differentiating between empty particles, spheroidal mRNA-loaded LNPs, and anisotropic multi-domain morphologies. Notably, AF4–SAXS validation of bleb-like morphologies displayed a distinct SAXS scattering signature in the pair distance distribution function, *p(r)*, showing that such unconventional structures can be identified beyond cryo-EM with detailed statistical and structural quantification. In addition, NanoFCM analysis revealed that the N/P ratio plays a critical role in defining effective particle loading, with LNPs formulated at N/P=6 exhibiting a more homogeneous mRNA content across the particle subpopulations. As such, we hypothesize that cores with a low degree of internal ordering combined with loading homogeneity could be key factors for the improved cell transfection observed at N/P=6. Altogether, these findings highlight how fractionation-based approaches can overcome the inherent complexity of LNP formulations, enabling the extraction of detailed structural features that are inaccessible to traditional ensemble-averaged methods and single-particle analyses.

## Results and Discussion

### Unveiling the Limitations of Ensemble-Averaged Methods

LNPs encapsulating 150 µg/mL (N/P=6) and 300 µg/mL (N/P=3) of GFP mRNA were formulated using a microfluidic device at identical lipid concentrations. DLS characterization of the LNP formulations displayed the expected single-exponential decay, yielding average hydrodynamic radius of 40.5 nm and 45.5 nm and polydispersity indexes of 0.09 and 0.07 for N/P=3 and 6, respectively (Fig. 1a). The intensity profiles and narrow distribution widths indicate the apparent presence of a single, relatively size-homogeneous LNP population (Fig. 1b). Further structural analysis of the unfractionated samples by SAXS suggests the presence of a relatively monodisperse population of slightly anisotropic spheroidal particles with a core-shell internal organization (Fig. 1c, Table S1). This is supported by the *ab initio* reconstruction of the LNP structure from the scattering profile, which provides a visual validation of this morphology. Thus, the analysis of the data from ensemble-averaged methods, i.e. DLS, SAXS, and NanoFCM, suggested the presence of a single particle population (Fig. 1d). However, cryo-EM characterization revealed clear structural heterogeneity, with spheroidal particles co-existing alongside unconventional morphologies, such as bleb-like structures and low-order aggregates (Fig. 1 e, Fig. S1).^8^ However, detecting them has been proved challenging with static methods, where particles at low concentrations marginally contribute to the signal and analysis methods are generally biased toward monomodal distributions.

**Fig. 1.**
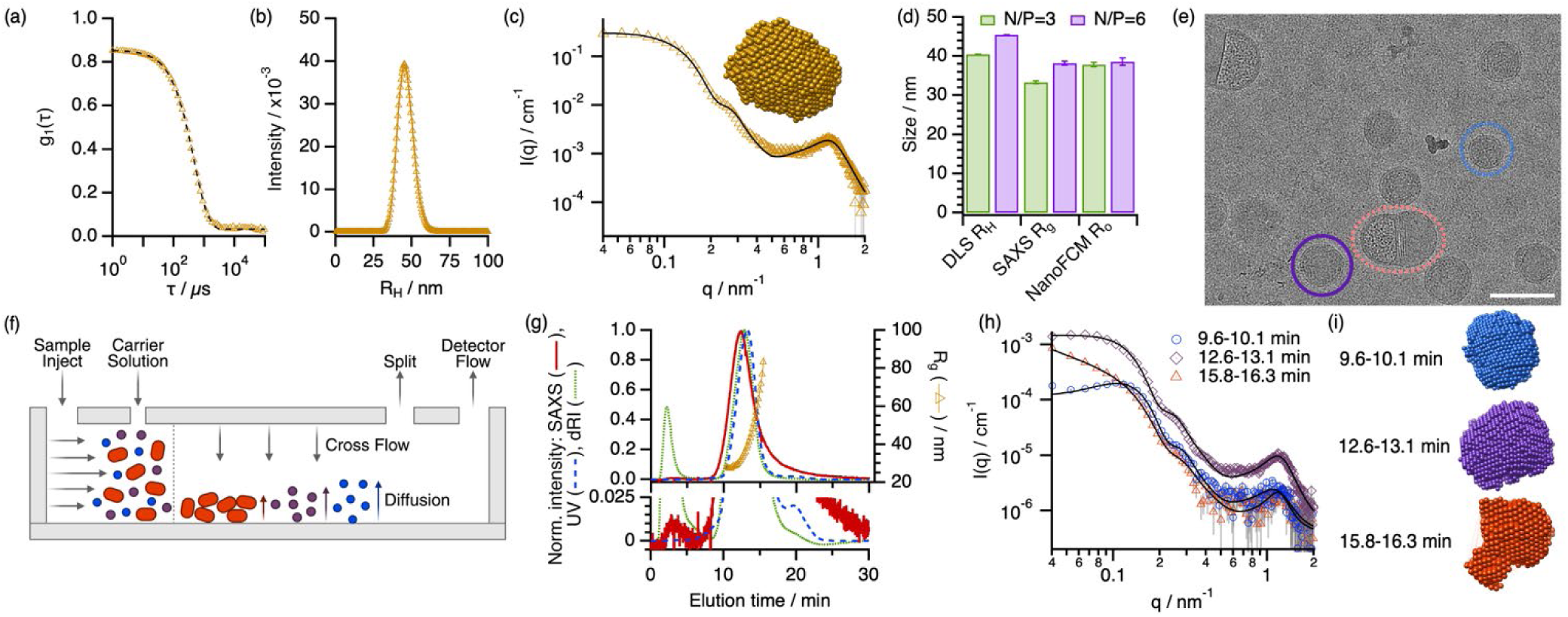
Overcoming Ensemble-Averaging in LNP Characterization Via AF4 Fractionation. (a) DLS correlogram, (b) calculated size distribution from (a), and (c) SAXS data (markers) and model (line) acquired for LNPs without prior fractionation. The inset in (c) shows the single-phase dummy atom model of the scattering profile. (d) LNP size obtained by different ensemble-averaged methods: hydrodynamic radius (*R_H_*) from DLS, radius of gyration (*R_g_*) from SAXS, and optical radius (*R_O_*) from NanoFCM. (e) Cryo-EM micrograph highlighting the presence of different structures. Scale bar = 100 nm. (f) Schematic representation of the AF4 configuration and function. (g) AF4 fractograms showing the evolution of scattering-corrected UV absorbance at 280 nm, differential refractive index (dRI), and solvent-subtracted integrated SAXS intensity with elution time. The right Y-axis displays the changes in *R_g_* as determined from multi-angle light scattering (MALS) coupled to the AF4 setup. (h) SAXS profiles (markers) and models (lines) from different fractions (as indicated in the legend of the graph) and (i) dummy atom reconstructions of the scattering profiles highlighting the morphological variety. Data were collected for a sample with a pre-injection concentration of 4.25 mM total lipid and 300 µg/ml mRNA (N/P=3). SAXS data were fitted using the core-shell ellipsoid model combined with the Teubner-Strey equation.^15^ Error bars represent the standard deviations from the averaged values; where not seen, error bars are within the markers.

To corroborate the cryo-EM findings that indicate particle heterogeneity, both N/P=3 and N/P=6 formulations were characterized by UV-vis spectroscopy, dRI measurements, LS, and SAXS in in-line coupling to AF4. AF4 fractionates polydisperse samples in the size range of 1-1000 nm in a single measurement under low shear forces in a channel without a stationary phase as required in chromatographic methods (e.g. size-exclusion chromatography, (SEC)).^33,34^ The diffusion coefficient of the particle—linked to its hydrodynamic size—serves as fractionation property and separation is achieved by use of minutely regulated laminar flows of carrier liquid (Fig. 1f). Compared to other fractionation techniques, such as SEC, AF4 exerts minimal interaction with the particles under separation. This makes AF4 particularly well suited for resolving distinct subpopulations of assembled systems, including LNPs, without altering their native structure. Scattering-corrected absorbance at 280 nm and dRI fractograms clearly showed an elution peak centered at ca. 13 min for both samples, suggesting a size-dependent fractionation of LNPs (Fig. 1g). In fact, the *R_g_* determined from MALS data confirmed a gradual increase in scatterer size within the elution window of this peak and reaches a 3-fold increase from the initial *R_g_* values. The results revealed that N/P=3 particles approximately showed double the absorbance area under the curve, consistent with its higher cargo concentration (Fig. S2). In addition, an early peak of lower intensity and width is observed at ca. 2.5 min. Considering the large dRI signal, the increase in the scattered intensity, and the absence of UV signal, this peak was attributed to the elution of formulation excipients, as confirmed by injecting a sucrose solution onto the AF4 system (Fig. S3). The absence of a UV peak arising from free mRNA is attributed to the high encapsulation efficiency (EE) of these LNPs, which have shown EEs of 92% for N/P=3 and 94% for N/P=6 in RiboGreen assays. This contrasts with previous AF4 characterization of mRNA-loaded liposomes that displayed a prominent pre-peak in the fractogram attributed to the elution of large amounts of free mRNA.^35^ Additionally, a shoulder in the UV signal was observed at ca. 20 min, possibly indicating a distinct subpopulation of larger LNPs with lower mRNA content, that became more prominent at the N/P ratio of 6 (Fig. S2).

Considering the elution profile observed in the previous fractograms, AF4-SAXS data were collected for both formulations for 35 minutes in 1 s frames (Table S2). SAXS detectors were placed in-line with the AF4 channel, with size-separated populations eluting and their scattering profiles directly analyzed. The AF4 setup used here was optimized for SAXS measurements by utilizing outlet flow splitting to prevent excessive dilution, enabling the collection of high signal-to-noise ratio even for low electron density samples, such as LNPs. In addition, tip sample injection with frit-inlet relaxation was utilized,^36^ which allows for higher injected sample mass without compromised fractionation compared to ordinary focus-relaxation channels.^37^ Thus, compared to previous investigations, ^19,35,38^ this configuration allows to study the overall and internal structure of LNP subpopulations. The evolution of solvent-subtracted integrated SAXS intensity with elution time showed a clear overlap with the UV and dRI signals (Fig. 1g), confirming that encapsulated mRNA is distributed across distinct particle subpopulations differing in size and volume fraction. To further investigate these differences, three representative fractions collected at different elution times were analyzed using a model-based approach combining a core–shell ellipsoid model with the Teubner–Strey equation (Fig. 1h). This combination of models enables the simultaneous description of the overall particle morphology and the internal structural ordering.^15,39^ A preliminary evaluation revealed an apparent increase in *R_g_* with elution time, from 23.1 ± 0.5 nm (9.6–10.1 min) to 30.4 ± 3.0 nm (12.6–13.1 min) and 76.7 ± 1.8 nm (15.8–16.3 min). This increase was attributed to the axial growth of prolate spheroidal particles, as discussed in the following section. Consistently, *ab initio* reconstructions of the scattering profiles displayed particles with progressively larger aspect ratios (Fig. 1i), in close correspondence with the morphologies observed by cryo-EM. These findings aligned with recent studies that suggested the presence of anisotropic particles co-existing with spheroidal assemblies.^24,25^ Together, these results highlight AF4–SAXS as a versatile platform for exploring the structural heterogeneity of LNPs, bridging the gap between ensemble-averaged and single-particle analyses.

### Characterization of Morphological Heterogeneity and LNP Sub-Population Distribution

To understand how structural diversity emerges within LNPs, we focused on the structure of distinct subpopulations across elution profiles isolated by AF4. AF4–SAXS data for N/P=3 and N/P=6 formulations within the main elution peak were divided into 13 fractions (Fig. 2a,b; Table S2) and analyzed using a combination of the Indirect Fourier Transformation (IFT) method and model-based fitting. The IFT analysis yields the pair-distance distribution function, *p(r)*, which describes the probability of finding two scattering points separated by a distance *r* within the particle, thereby providing real-space information about particle size and shape.^40,41^ The *p(r)* profiles were calculated for each AF4 fraction and normalized to the scattered intensity at zero angle (*I(0)*), *p(r) I(0)^-1^*), revealing similar trends for both N/P ratios (Fig. 2c,d). Fractions eluting between approximately 10 and 14 min exhibited relatively symmetric *p(r)* curves, consistent with particles of low anisotropy. In contrast, later fractions displayed asymmetric *p(r)* profiles with pronounced tails that evolved into a secondary maximum, of which its intensity and maximum dimension increased with elution time (Fig. S5). In SAXS analysis, an asymmetric or bimodal *p(r)* is indicative of elongated or multi-domain structures, whereas symmetric profiles correspond to isotropic particles.^42^ Therefore, this progressive development of a secondary peak in the *p(r)*, coinciding with later eluting fractions—and thus larger particles— confirmed that the increase in particle size is associated with a transition from near-spherical to anisotropic morphologies.

**Fig. 2.**
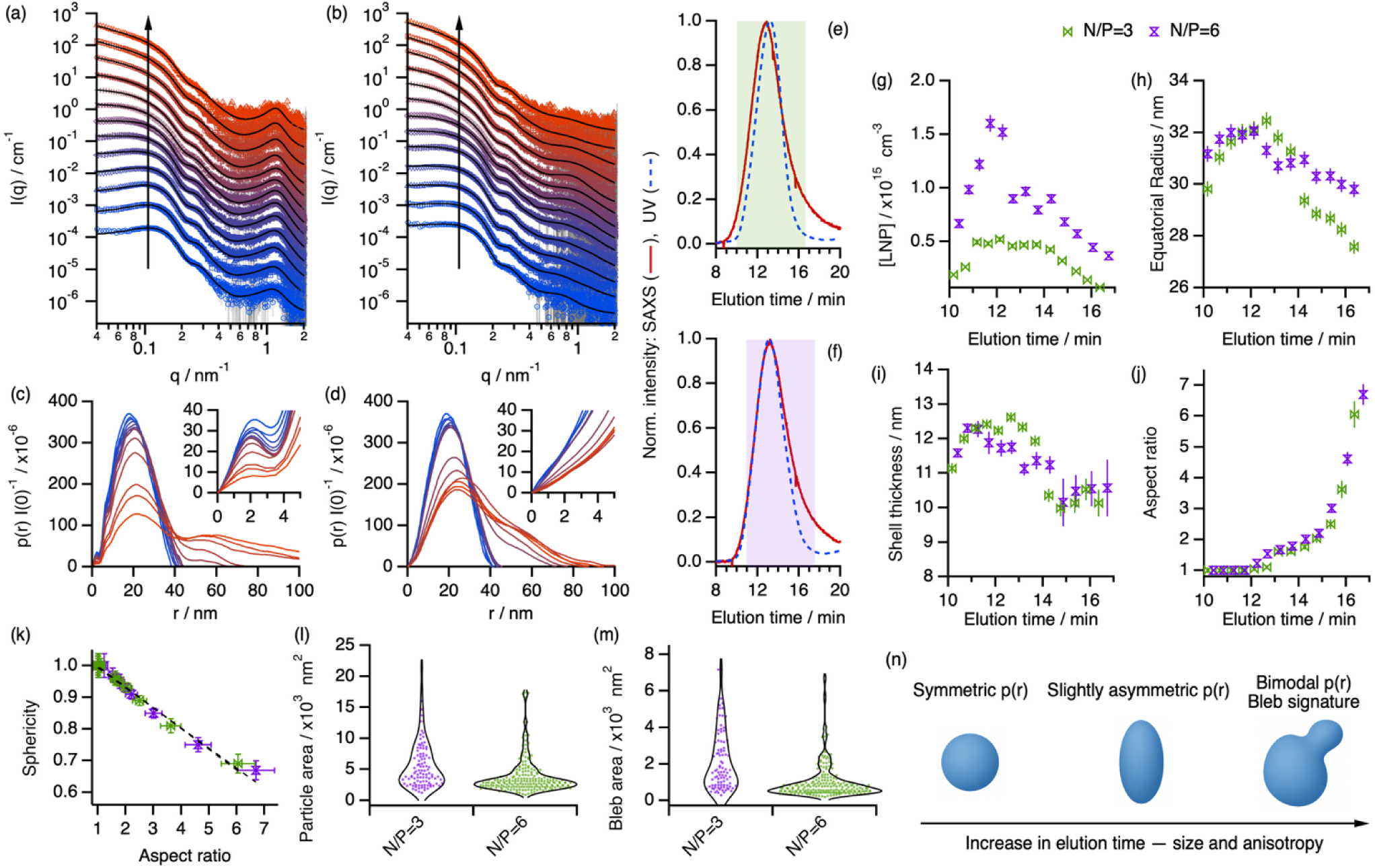
AF4–SAXS Characterization of LNP Subpopulations Highlights Size-Dependent Morphological Evolution. AF4-SAXS data (markers) and fits (black lines) form model-based analysis for (a) N/P=3 and (b) N/P=6 nanoparticles. Data have been offset for clarity and elution time increases in the direction of the arrows. (c) and (d) show the *p(r)* profiles derived from IFT data analysis for the data shown in (a) and (b), respectively. AF4 fractograms showing the evolution of the normalized scattering-corrected UV absorbance at 280 nm and solvent-subtracted integrated SAXS intensity with elution time for (e) N/P=3 and (f) N/P=6. Shaded regions indicate the portions of the AF4 elution peak that were binned in 30 s intervals for subsequent SAXS data analysis. Main results from the structural characterization by AF4-SAXS: (g) particle concentration, (h) equatorial radius, (i) shell thickness, and (j) aspect ratio. (k) Parametric plot showing the correlation between particle sphericity and aspect ratio. Results from the analysis of cryo-EM micrographs: projected (l) particle area and (m) bleb area, defined as the area occupied by blebs within a particle. Particles analysis was performed using Fiji.^43^ Data were collected for samples with a pre-injection concentration of 4.25 mM total lipid and 150 µg/ml (N/P=6) or 300 µg/ml mRNA (N/P=3). Model-based analysis of SAXS data was performed using the core-shell ellipsoid model combined with the Teubner-Strey equation.^15^ Error bars represent the standard deviations from the averaged values; where not seen, error bars are within the markers. (n) Schematic representation of LNP structural evolution across the AF4 elution profile: early fractions contained compact spherical particles, intermediate fractions related slightly anisotropic prolate spheroids, and later fractions exhibited highly anisotropic, bleb-like assemblies.

This compositional heterogeneity suggests the need to couple fractionation with model-based analysis of the AF4-SAXS data to further investigate particle structure. Although most SAXS and SANS studies of LNPs have employed core–shell spherical models,^15,44^ and some have extended to two-compartment cylinder models to reflect more complex morphologies observed by cryo-EM,^13^ there remains a lack of systematic modelling addressing LNP anisotropy across subpopulations. Here, we employed a core–shell prolate ellipsoid model to describe the overall particle geometry within the different fractions.^45^ This model can approximate the scattering of spherical LNPs when the aspect ratio (*AR = r_po_/r_eq_*, where *r_po_* and *r_eq_* are the polar and equatorial radii, respectively) approaches unity, capture subtle deviations in particle shape when *AR* > 1, and describe highly anisotropic particles when *AR* ≫ 1. In addition, other parameters, such as particle concentration and scattering length density distribution within the particle (Table S3), can be extracted from the analysis.

The evolution of the UV and integrated SAXS intensity with elution time suggests a non-monotonic transition in particle concentration across the AF4 separation, where the maxima in the scattering-corrected absorbance at 280 nm is observed at 13.1 min and 13.3 min for N/P=3 and N/P=6 formulations, respectively (Fig. 2e,f). From the analysis of SAXS data we observed that particle concentration, parametrized as the number of particles per unit volume (Fig. 2g), was relatively low at early elution times (< 12 min), followed by a pronounced increase that reached a maximum at intermediate elution times (≈ 13 min) before decreasing again toward later fractions (> 14 min). The observed non-monotonic profile reflects the coexistence of LNP subpopulations differing in concentration and morphology, consistent with the kinetic trapping and polydisperse nature of LNP self-assembly during microfluidic mixing.^25,46,47^ Thus, the concentration maximum observed at intermediate elution times coincides with the more prevalent spheroidal LNPs as these display a symmetric *p(r)*. In contrast, later fractions with lower particle concentration are enriched in anisotropic morphologies. The main population at these intermediate times displayed a 4-fold higher concentration for N/P=6 and a seven-fold higher concentration for N/P=3 compared to the late-eluting fractions. The proportion of anisotropic particles was determined by integrating the particle density across AF4 fractions exhibiting asymmetric *p(r)* distributions. This analysis revealed that anisotropic particles accounted for 30 ± 2 % of the total population in the N/P=3 formulation and 23 ± 2 % for the N/P=6 ratio, suggesting that lower N/P ratios promote increased morphological heterogeneity for our formulations. Notably, LNPs at N/P=6 consistently exhibited higher overall particle concentrations throughout the elution window, although this trend may partially reflect systematic variations during sample preparation or characterization.

The overall radius of the particle (core + shell) from SAXS measurements ranged between 30 nm and 32 nm for spheroidal particles (elution time < 14 min, Fig. 2h). This overall dimension can be divided in two regions with different scattering length densities (SLDs), a shell and a core with low and high electron density, respectively. It should be noted that, due to the constraints of the SAXS model employed, the particle core is treated as a uniform region enclosed by the shell, and therefore encompasses both the ionizable-lipid–rich interior and any bleb-like protrusions, without distinguishing between these domains. The shell thickness remained relatively unchanged across the different particle subpopulations and N/P ratio (Fig. 2i), displaying an average thickness of 11.2 nm. Notably, the low SLD values observed for the shell region (≈ 8.4×10^-6^ Å^-2^) for both N/P=3 and 6 suggested that the shell was formed by a lipid bilayer (Table S3). As unilamellar shells adopt thicknesses around 5 nm,^46^ the differences in the shell dimensions are likely attributed to the presence of solvated PEG chains.^11^ The core presented higher SLD than that of the solvent (SLD_buffer_ = 9.4×10^-6^ Å^-2^), with values ranging between 11×10^-6^ Å^-2^ and 12×10^-6^ Å^-2^. Considering the low SLD of ionizable lipids, this significant increase in the SLD was attributed to the localization of mRNA at the LNP core.^8,46^ Notably, changes in the core SLD with elution time suggested different particle loading across LNP subpopulations, which will be discussed in the following section.

Further analysis revealed that the apparent increase in particle *R_g_* observed from AF4-MALS results and IFT analysis primarily emerged from LNP elongation along the polar axis (Fig. 2j). At early elution times (<12 min), particles exhibited near-spherical morphology (*AR* ≈ 1.0), which gradually transitioned to mildly anisotropic shapes up to 15 min (*AR* ≈ 1.6). Beyond this point, a sharp increase in aspect ratio indicated pronounced anisotropy, likely associated with the elution of unconventional structures such as bleb-like morphologies and low-order aggregates (*AR* > 1.6). It should be noted that particles with the largest AR values likely corresponded to the presence of fused/aggregated LNPs, but the limited q-range of the experiment did not allow to resolve the structure of those. Accordingly, the second oscillation observed in the *p(r)* distributions can be attributed to the presence of bleb-like domains at the particle surface. To the best of our knowledge, this represents the first demonstration of an alternative method to cryo-EM for the detection, quantification, and structural characterization of bleb-like features in LNP formulations. When comparing LNP formulations at N/P=3 and N/P=6, subtle differences emerged in the particles’ aspect ratios (Fig. 2j). At early elution times, the two formulations displayed similar morphology. At intermediate elution times—where particle concentration peaked—the N/P=6 particles were slightly larger, which likely explains why the static characterization methods (DLS, light scattering, SAXS) reported larger apparent diameters for the N/P=6 series. At later elution times, however, N/P=3 particles became larger as they adopted more anisotropic shapes, consistent with a higher prevalence of elongated or bleb-like morphologies compared to the formulation with higher lipid content. In fact, a closer look at the sphericity parameter revealed a higher tendency of N/P=3 particles to form larger and more anisotropic assemblies than the N/P=6 formulation (Fig. 2k), thus directly correlating particle asymmetry to the preferential growth in the polar dimension. This suggests that a lower N/P ratio (higher mRNA loading relative to lipid) drives greater morphological polymorphism and shape heterogeneity, in agreement with previous reports.^28^

The results from the analysis of AF4-SAXS data were corroborated by cryo-EM micrograph analysis of each unfractionated sample (Fig. S6). While the analysis of SAXS data could not provide direct information of bleb dimensions, cryo-EM analysis can reveal particular details about the characteristics of the LNPs, such as the projected particle area and bleb area (here referred to as the area occupied by a bleb for a given particle). For both N/P=3 and 6, the violin plots of the particle’s projected area showed the presence of small particles with projected areas around 3000 nm^2^ with higher statistical frequency than unconventional morphologies associated with larger areas (Fig. 2l). N/P=3 particles displayed a broad distribution skewed toward larger areas, consistent with the asymmetry observed by the analysis of SAXS data. In contrast, the narrower, smaller-area distribution for N/P=6 indicated reduced morphological diversity. By analyzing the areas occupied by the blebs, we observed that N/P=3 formulations also indicated a broader area distribution, suggesting that domain segregation and bleb-like morphologies are critical factors for defining overall particle size. On the other hand, LNPs formulated at N/P=6 primarily showed the formation of regular blebs with projected areas around 700 nm^2^. Therefore, our combined analyses clearly demonstrated that formulations at lower N/P value yield higher structural variability in the LNPs, which is associated with higher prevalence of particles with one or multiple blebs (Fig. S5).

### Internal Architecture and Cargo Loading Across Subpopulations

Having established the presence of distinct LNP subpopulations differing in size and morphology, we next examined their internal architecture to understand how structural variations relate to mRNA loading. AF4-SAXS data of individual fractions provided insight into differences in internal organization, showing how mRNA content and structural ordering evolve across the elution profile as a function of the N/P ratio. Visual observation of the SAXS data consistently revealed a distinct peak centered at *q*≈1.2 nm^-1^ for the N/P=3 system that was absent for the formulation with higher lipid content (Fig. 3a,b). As in any diffraction-based method, the presence of such a peak indicates the existence of ordered domains, confirming the formation of more ordered particle cores for N/P=3 particles compared to the disordered cores for the N/P=6 system. Cryo-EM micrographs further corroborate this observation, displaying the characteristic oscillatory texture associated with quasi-periodic internal organization for N/P=3 particles (Fig. 3c). In contrast, N/P=6 particles showed a more homogeneous intensity distribution (Fig. 3d). As expected, gray-scale intensity values across the particle cores indicate higher electron density in N/P=3 particles (Fig. 3e), attributed to a greater mRNA accumulation relative to the lipid. This behavior has been previously attributed to the stronger electrostatic interactions and tighter packing between mRNA molecules and ionizable lipids within the particle core, which becomes more prominent at higher mRNA contents.^29^ Consistent with this, the corresponding *p(r)’s* displayed a pronounced short-range oscillation centered at ≈2.5 nm in the N/P=3 system that was not observed at the higher lipid content (Fig. 2c,d).

**Fig. 3.**
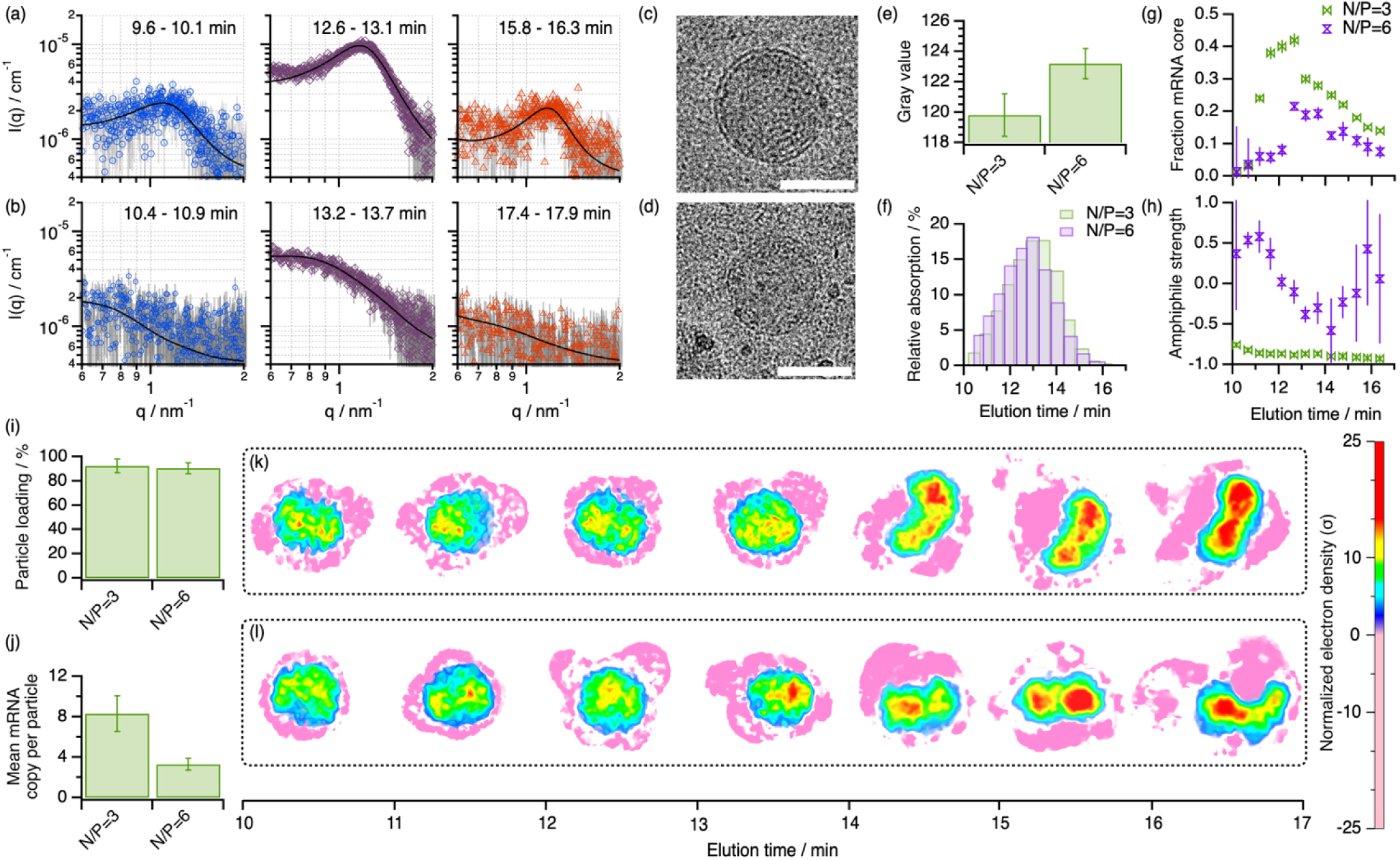
Evolution of Internal Ordering and Domain Segregation Across LNP Subpopulations. AF4-SAXS data (markers) and models (solid lines) showing the evolution of the high q expansion of the data for particles formulated at N/P (a) 3 and (b) 6 at different elution times (as indicated in the legend of each graph). Micrographs of LNPs at N/P (c) 3 and (d) 6 focused on the structure of a representative particle. Scale bars = 50 nm. (e) Gray values extracted from the LNP micrographs shown in (c) and (d). Background gray value was 130±1. (f) Evolution of the scattering-corrected relative absorbance at 280 nm for N/P=3 and N/P=6. (g) Fraction of the particle core that is occupied by mRNA as determined from SAXS data analysis. (h) Amphiphile strength calculated from the results of the Teubner-Strey modelling. Results derived from NanoFCM analysis: (i) percentage of particles containing at least one mRNA copy and (j) average mRNA copy per particle. Particle electron density reconstructions from the SAXS data using DENSity from Solution Scattering (DENSS) for LNPs formulated at (k) N/P=3 and (l) N/P=6.^50^ The relative electron density shows how many standard deviations a region’s density differs from the average solvent background. Electron dense regions correspond to mRNA-rich and negative density corresponds to hydrophobic lipid regions. Error bars represent the standard deviations from the averaged values; where not seen, error bars are within the markers.

These findings confirmed a clear link between internal ordering and mRNA loading, motivating a closer examination of how periodicity evolves with elution time. Analysis of the AF4-SAXS data revealed significant changes in both shape and intensity across the elution profile, suggesting that the degree of internal ordering also evolved with fractionation. Initially, we wanted to confirm whether these changes could be correlated to the amount of mRNA at the particle core. The absorbance data at 280 nm showed that the mRNA content within each fraction varied non-monotonically, reaching a maximum at intermediate elution times and coinciding with the elution range of the most concentrated particle populations (Fig. 3f). To decouple concentration effects, we used the fitted scattering length density of the particle core to estimate the mRNA content per particle. The results showed a similar non-monotonic trend; individual particles at early elution times contained relatively low amounts of mRNA, followed by a maximum loading at intermediate times, and a subsequent decrease at later elution times (Fig. 3g). As expected, the N/P=6 formulation exhibited a lower overall mRNA content (< 20% of the core volume) than the N/P=3 system, while also showing less pronounced variations in loading across fractions. These findings were further corroborated by NanoFCM measurements (Fig. S7). By differentiating between fluorescence-positive from fluorescence-negative particles in the presence of the mRNA-tagging fluorescent dye SYTO-9, we could determine the number of particles with mRNA at the core and compare that to the number of empty particles. Our results confirmed that more than 90% of the total particle population contained mRNA (Fig. 3i),^48^ contrasting with other formulation protocols that suggested that only around 60% to 70 % of the particles were loaded with mRNA.^25,49^ Considering the mRNA content in each fraction, early-eluting LNPs could comprise the majority of unloaded particles. Moreover, N/P=3 formulations displayed a higher average mRNA content per particle (Fig. 3j), as expected from the lower amount of ionizable lipid relative to the mRNA content. However, this is accompanied by greater variability, consistent with an increased heterogeneity in mRNA encapsulation between subpopulations

We next investigated how mRNA is organized within the particle core. To quantify internal ordering, SAXS data at high *q* were analyzed using the Teubner–Strey model, which describes systems with quasi-periodic electron density fluctuations.^39^ This model defines two structural parameters: the periodicity length (*d*), which reflects the average spacing between electron density fluctuations (e.g., lipid–mRNA domains), and the correlation length (*ε*), which measures how far this local order extends before becoming random. The analysis revealed that both structural parameters remained relatively constant across subpopulations for the N/P=3 system, with an average periodicity length of ≈5.3 nm and correlation length of ≈3.1 nm. As the periodicity length corresponds to the center-to-center repeat distance between regions of high electron density, the repeating motif likely consisted of lipid bilayer domains separated by mRNA-rich layers. However, the shorter average correlation length indicated only short-range order rather than extended multilamellar stacks. These results suggest the presence of a percolating quasi-lamellar lipid domains templated by the presence of mRNA. This model agrees with recent coarse-grained MD simulations of LNPs at similar N/P ratios that suggest the presence of bicontinuous lipid-mRNA domains surrounded by water channels that limit orientational periodicity.^51^ In addition, the absence of significant changes in internal ordering across the N/P=3 fractions suggested that mRNA encapsulation possibly reached a saturation threshold,^28^ where the particle core is primarily defined by charge-neutralized mRNA–lipid complexes. Beyond this, additional mRNA cannot be accommodated without disrupting the overall particle architecture or altering functional loading.^52^ In contrast, at higher N/P ratios the diffraction peak disappeared, indicating the loss of long-range ordering and a transition to a more disordered core structure. In this case, the variations in *d* and *ε* across the elution profile suggested changes in the local organization of mRNA-lipid complexes, again reflecting the differential loading observed across subpopulations.

Using the parameters derived from the Teubner-Strey model we computed the amphiphile strength, an empirical dimensionless quantity that reflects the degree of internal ordering and the type of quasi-periodic structure present.^39^ For the N/P=3 system, the value evolved from -0.86 at early elution times to almost -1 for later fractions (Fig. 3h), indicating a higher order at the LNP core with increasing mRNA content. Notably, this value of -1 is commonly attributed to the presence of lamellar structures, which is the predominant ordered phase in mRNA-loaded LNPs.^7,15^ Conversely, the degree of ordering for N/P=6 particles changed non-monotonically, where the higher degree of ordering—reaching values close to -0.6 for intermediate elution times—overlapped with the mRNA loading maximum. At early (<12 min) and late (>15 min) elution times, the amphiphile strength became positive, a signature of highly disordered internal structures.^53^

These results demonstrated that internal order within LNPs is tightly coupled to mRNA loading and formulation stoichiometry. Building on our previous observation of anisotropic particles within the formulations, we next examined how structural anisotropy elution times influences the internal structure of the particles. The well-defined peak observed in the N/P=3 formulation allowed us to further investigate mRNA ordering at the particle core employing a peak analysis approach (Fig. 3a). For most elutions, deconvoluting each peak in our data with two Lorentzian components significantly improved the quality of the fits compared to the use of a single Lorentzian component (Fig. S7), suggesting the presence of a second ordered phase at pH 7.4.^54^ A high-q peak corresponding to average d-spacings of ≈ 5.0 nm was observed at all elution times, while a low-q peak attributed to a d-spacing of ≈ 5.8 nm gradually appeared at late elutions (Table S4). Recent studies have shown the presence of multiple particle subpopulations by deconvoluting asymmetric peaks into two peaks.^17^ However, as our approach was already fractionating particles by size, we hypothesize that this dual-peak instead depicted the coexistence of distinct lipid-mRNA domains within individual particles. Notably, this becomes more prominent in fractions that are attributed to bleb-like morphologies.^8^ To further characterize internal structure, the internal electron density distributions of the LNPs were reconstructed using DENSS.^50^ For both N/P ratios, the pink regions of negative contrast (electron density <0 relative to the solvent) corresponded to the low electron-density regions formed by the lipid shell, while the red and yellow electron-rich domains at the core contained the mRNA-rich regions complexed with ionizable lipid (Fig. 3k,l). Intermediate blue/green areas represent medium density regions that may contain averaged heterogeneous internal structures of ionizable lipid-RNA complexes with lower mRNA density. This aligns with previous studies that reported the co-existence of complexes with different mRNA densities at the LNP core.^25,54^ In terms of overall structure, at early elution times (< 14 min), LNPs showed spheroidal geometries with a single mRNA-rich core (Fig. S8, S9). However, at later eluting fractions, LNPs developed overall anisotropy accompanied by a clear segregation of electron-dense regions, forming two discrete mRNA domains separated by a thin lipid layer of lower electron density. This transition from single-core to multi-domain internal architectures confirms the formation of bleb-like structures within anisotropic particles. Notably, the elution range where domain segregation appears overlapped with the transitions observed by IFT and model-based analyses, providing unequivocal evidence for the coexistence of multiple particle populations as validated across orthogonal AF4–SAXS analytical approaches and cryo-EM.

### Structure-function relationship between LNP morphology and in vitro performance

To study the implications of the observed structural heterogeneity in LNP function, we next evaluated the in vitro transfection efficiency of our formulations. The goal was to shed some light on how differences in internal architecture and cargo distribution translate into delivery efficacy. As demonstrated in the preceding sections, N/P=3 formulations exhibit more ordered internal structures and a higher degree of mRNA loading, whereas N/P=6 particles are less ordered but display greater morphological uniformity. Given these distinctions, we hypothesized that the balance between internal ordering, anisotropy, and particle loading would directly influence endosomal escape mRNA release and protein expression. To test this, dose–response transfection assays were performed in HEK293 cell lines under equivalent mRNA concentrations (Fig. 4a). As expected, the expression of eGFP after the internalization of the encoding mRNA was observed, where higher doses of mRNA yielded increased protein expression. From those signals, the mean fluorescence intensity was determined and plotted as a function of mRNA dose and N/P ratio (Fig. 4b). We observed that the N/P=6 formulation consistently displayed higher transfection efficacies than the N/P=3 counterpart. For a constant mRNA dose, increasing the lipid content—thus N/P ratio—often results in improved cytosolic delivery while keeping the mRNA concentration constant.^3,28^ When comparing equivalent lipid concentrations at different mRNA doses, e.g. 50 ng mRNA for the N/P=6 formulation used the same amount of lipid as the 100 ng mRNA for the N/P=3 formulation, we can isolate the payload loading effect from the lipid concentration effect. Our results confirmed that the N/P=6 formulation improved protein expression by at least 25% despite the higher mRNA doses in the N/P=3 system (Fig. 4c). Only the lowest lipid dose showed no detectable difference between formulations (p ≈ 1), likely reflecting the overall low transfection levels under these conditions and masking formulation-dependent effects.

**Fig. 4.**
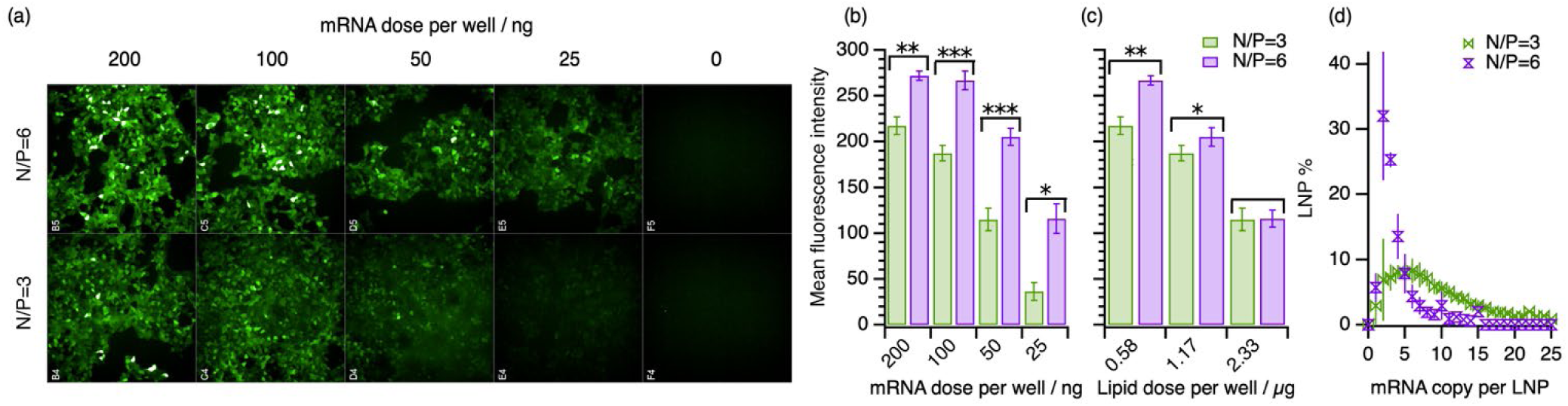
Correlation Between In Vitro Transfection Efficiency, Structural Heterogeneity, and Cargo Loading. (a) Representative confocal images of HEK293 cells subjected to varying doses of mRNA for the two N/P ratios. Micrographs in the last row correspond to the control experiments in the absence of mRNA. Experiments were performed in triplicates. Transfection efficacy parametrized as the mean fluorescence intensity of expressed eGFP for the experiments shown in (a) for different (b) mRNA or (c) total lipid doses. Data are shown for different mRNA doses using either the N/P=3 or 6 formulation, as indicated in the legend of the graph. The values show the average mean fluorescence intensity for each condition and the error bars correspond to the standard deviations for the triplicates. Significance was assessed by Two-way ANOVA, with \**p* < 0.05, \*\**p* < 0.01, \*\*\**p* < 0.001. Comparisons with *p* ≥ 0.05 are left unmarked. (c) Loading distribution profiles as a function of mRNA copy per particle for N/P=3 and 6, as indicated in the legend of the graph. The values from NanoFCM experiments correspond to the averages of 4 independent experiments and error bars display the standard deviation from the averages.

We next sought to identify the structural features responsible for the enhanced transfection performance of the N/P=6 formulation. In particular, we examined how differences in particle loading correlated to the delivery efficacy. Single-particle NanoFCM analysis provided quantitative information on LNP loading across individual LNPs, revealing distinct loading distributions between formulations. The N/P=3 formulation exhibited a broad loading profile, with particles encapsulating between 1 and 21 mRNA copies and a subtle maximum at 6 copies, corresponding to approximately 6% of the total population. It should be noted that only a 2% of particles displayed >17 mRNA copies, which could possibly be attributed to the presence of more than one particle or a fused aggregated at the detection pathway. In contrast, the N/P=6 system displayed a narrower, well-defined distribution centered around 2 mRNA copies per particle, accounting for roughly 32% of the total number of LNPs. Notably, mRNA copy numbers between 2 and 4 corresponded to more than 70% of the particles, highlighting the improved homogeneity of the N/P=6 formulation. This made us hypothesize that this homogeneous loading, together with the presence of a less ordered particle core and reduced particle heterogeneity, likely contributed to the improved transfection efficacy of the N/P=6 system compared to the formulation at higher mRNA content. Overall, these results underscore that structural and loading uniformity enhance delivery performance, disentangling compositional effects from those arising from internal organization.

## Discussion and Outlook

In this work, we challenged the traditional approaches for the characterization of mRNA-loaded LNPs. Conventional methods have long struggled to reconcile two critical aspects of LNP analysis: the need for detailed structural information and the requirement for statistically meaningful data. Techniques such as cryo-EM provide high-resolution snapshots of individual particles but cannot capture ensemble diversity,^18^ whereas ensemble-averaging methods like DLS lack size and morphological resolution and SAXS obscures the contributions of low-abundance particles.^17^ Yet, unusual assemblies—such as bleb-like structures—are regarded as functionally relevant subpopulations for transfection efficiency.^16^ While the development of holistic characterization strategies has recently gained significant attention,^19,25^ detailed characterization of both external structure, internal organization, and loading profile for coexisting particle subpopulations has remained elusive. The integrated AF4–UV-MALS-dRI-SAXS and NanoFCM framework presented here overcomes these limitations by combining size-based fractionation, structural reconstruction, and single-particle quantification to resolve both the overall architecture and loading heterogeneity of LNPs with statistical robustness.

Our multidimensional approach revealed that LNP formulations comprise structurally distinct subpopulations. Size-based separation by AF4 combined with SAXS facilitated optimal structural characterization of LNP samples,^19^ enabling phase identification, mRNA distribution, and the elucidation of morphological variants across particle subpopulations. This allowed us to develop detailed structural models from SAXS data by combining IFT analysis, model-based fitting, and DENSS reconstructions. Our characterization revealed that the main particle population can be described as prolate spheroidal assemblies with subtle anisotropy. Also, the presence of significant amounts of anisotropic assemblies (> 20%) with significant asymmetry was identified. In fact, our strategy provides a new approach to detect and quantify the presence of unconventional assemblies, such as bleb-like morphologies and low-order aggregates, allowing isolation of these structures for improved structural resolution and statistical validation compared to single-particle analysis using cryo-EM. In addition, our findings established a clear link between LNP overall heterogeneity and internal architecture. We showed how mRNA preferably sits at the core of spheroidal assemblies, while small particles and anisotropic morphologies (e.g. blebs) effectively encapsulate lower amounts of payload relative to the lipid content. Therefore, AF4-coupling to advanced characterization methods, such as SAXS, provides a powerful approach to extract structural information without perturbing the native characteristics of the system due to interactions with the stationary phases used in common SEC chromatographic methods.^55^

The combination of NanoFCM with AF4–SAXS provides a powerful framework for establishing direct structure–loading relationships in complex LNP systems. Together, these methods bridge the gap between ensemble-averaged and single-particle analyses, providing complementary information that neither technique can deliver alone. While AF4–SAXS enables quantitative assessment of morphological and internal structural features across size-resolved subpopulations, NanoFCM adds single-particle statistical precision by quantifying cargo content. This dual analytical approach gains particular relevance when investigating formulations at different N/P ratios. Critically, we could recognize loading homogeneity and disordered particle cores as key determinants for the improved delivery efficacy observed for the N/P=6 formulation, suggesting that molecular mobility may facilitate cargo release following cellular uptake.^30^ In contrast, the more ordered cores and higher loading heterogeneity observed at lower N/P ratios likely hinder efficient unpacking and translation, despite higher overall mRNA content. Therefore, controlling the balance between structural stability, cargo distribution in LNPs, and dynamic disorder of nucleic acid-lipid complexes emerges as a key parameter for optimizing both *in vitro* potency and, potentially, *in vivo* performance. These insights underscore the importance of integrating structural characterization with functional assays to inform predictive design rules for next-generation LNP-based RNA therapeutics.

Overall, our multi-scale approach enables a holistic and statistically meaningful description of LNP populations, advancing the development of predictive models for structure-dependent delivery efficacy, including endosomal escape and biodistribution. The combination of AF4 with in-house detection (e.g. UV-vis, dRI, MALS, DLS) can open an easy-to-access route to assess particle heterogeneity. In addition, the integration of those methods with advanced characterization, AF4-SAXS and NanoFCM, can bridge population-level quantification with high-resolution structural analysis, which are gradually becoming more widely available due to the user-friendliness of analysis protocols.^41,56^ By strengthening the ability to link structure to functional outcomes, a holistic approach is critical for advancing design and delivery of LNP therapeutics.

## Methods

### Formulation of LNPs

LNPs were prepared in 20 mM Tris-HCl buffer, and consists of cationic ionizable lipid, 7-[(2-Hydroxyethyl)[8-(nonyloxy)-8oxooctyl]amino]heptyl 2-octyldecanoate (Lipid 5, 99.28%, MedChemExpress), cholesterol (> 99%, Sigma Aldrich), distearoylphosphatidylcholine (DSPC, > 99%, Avanti Polar Lipids) and 1,2-Dimyristoyl-snglycero-3-methoxypolyethylene glycol (DMG-PEG2k, > 99%, Avanti Polar Lipids). The samples consisted of encapsulated GFP mRNA formulated at N/P ratios 6 and 3. Lipids were dissolved in ethanol at molar ratios of 50:10:38.5:1.5 (lipid 5:DSPC:cholesterol:DMG-PEG2k) at a total lipid molar concentration of 4.25 mM. The lipid mixture was combined with 0.2 or 0.4 mg/mL GFP mRNA (CleanCap eGFP, TriLink) in 25 mM acetate buffer (pH 4.5) at a ratio of 3:1 vol/vol (aqueous:organic) and at a total flow rate of 12 mL/min using a microfluidic mixer (NanoAssemblr® Ignite™, Precision NanoSystems) to obtain N/P ratios of 6 or 3. The LNP formulations were subsequently dialyzed in Slide-A-Lyzer G3 cassettes (10 k MWCO) against 20 mM Tris-HCl Buffer (pH 7.4) at 5 °C for 1 day with a fresh buffer exchange after 1 h. Sucrose was added to reach 8% (w/w) and the LNPs were concentrated using Amicon Ultra 4mL filters. Finally, samples were adjusted to final concentrations of 300 and 150 µg/mL mRNA for N/P=3 and 6, respectively, and a total lipid concentration of approximately 3.5 mg/mL. mRNA concentration and encapsulation efficiency (EE) were evaluated via Ribogreen assay according to the manufacturer’s protocol.

### Dynamic Light Scattering

DLS measurements were performed using the DynaPro® Plate Reader III (Wyatt Technology, Santa Barbara, CA, USA). Samples were loaded into a 96-well clear-bottom polystyrene microplate (Corning 3540), sealed to prevent evaporation, and equilibrated at the desired temperature prior to measurement. The samples were diluted 50 times in PBS. Measurements were conducted at 25 °C using DYNAMICS® software (Wyatt Technology) for data acquisition. Each sample was measured in triplicate with an acquisition time of 5 seconds per read and 40 reads per well. Intensity autocorrelation functions were averaged and analyzed using Igor Pro® 8.04 using the cumulant equation—with the first cumulant representing the average decay rate and the second cumulant reporting on the variance of the decay—to determine the hydrodynamic radius, intensity distribution, and the polydispersity index.

### Asymmetric Flow Field-Flow Fractionation

The samples were fractionated on an AF4 system from Waters-Wyatt Technology Europe GmbH (Germany), equipped with an Eclipse Neon for flow control, with dilution control module (DCM) for split-outlet application, and a dispersion inlet (DI) channel for stop-less tip-injection frit-inlet application, with a 275 µm thick, wide (W) spacer, and a 10 kDa ultrafiltration membrane from regenerated cellulose acting as accumulation wall. Optilab T-rEX and Dawn Heleos II (Wyatt Technology, USA) were used for acquisition of refractive index (RI) and multi-angle light scattering (MALS) signals respectively. A UV detector from Agilent was used for acquisition of UV absorption/scattering at wavelength 280 and 340 nm. As a final detector, the eluting fractions were directed to the in-line coupled SAXS capillary of the BioCube at the CoSAXS beamline, MAX IV, Sweden. The carrier liquid flow was delivered by an isocratic pump, and the samples were injected using an autosampler, both from Agilent Technologies 1100-series (USA). The carrier liquid was a Tris-HCL buffer (25 mM, pH 7.4) containing 50 mM NaCl.

The fractionation method started at a constant cross flow of 2.5 ml/min for 5 min, followed by a linear decaying cross flow to 0.3 ml/min with a half-time of 3 min, followed by a linear decaying cross flow to 0.05 ml/min with a half-time of 10 min. The channel flow was 1.75 ml/min and the detector flow 0.2 ml/min. The injection flow was constant throughout the method, at 0.2 ml/min.

To maximize the concentration in the eluting fractions, the detector flow was minimized—i.e. split-outlet flow maximized—and the injected sample volume was 200 µl with an approximate concentration of 3.5 µg/µl based on lipids, optimized by increasing injection volume without any sign of channel overloading or significant peak broadening. The *R*_g_ values were determined from MALS data. To avoid distorted *R*_g_ values, Debye 1st order fitting was performed on two flanking intervals before and after the elution peak maximum. *R*_g_ values at the peak maximum were interpolated from these fits.

### AF4-UV dual wavelength correction

To quantify mRNA content across the LNP size distribution, an AF4-UV dual wavelength method was employed. At 280 nm, the signal intensity, *I*, comprises both mRNA absorbance and particle scattering. To disentangle these contributions, simultaneous measurements at wavelengths *λ*=280 nm and *λ*=340 nm were taken. Since UV absorbance at 340 nm is negligible, the intensity at this wavelength is dominated by scattering and can be converted by a conversion factor, *k*, to its equivalent at 280 nm using the following relationship:

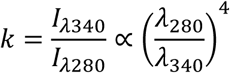

This correction leverages Rayleigh scattering principles, applicable when particle sizes are smaller than the incident light wavelength, where the Rayleigh ratio is proportional to *λ*^-4^. Given that the molar extinction coefficients of single- and double-stranded mRNA are comparable at 280 nm, this wavelength was deemed appropriate for absorbance measurements within the Rayleigh regime. Departing from the conventional wavelength used for RNA absorption, 255-260 nm, was made to minimize error in the Rayleigh approximation.

By subtracting the converted scattering signal from the total signal at 280 nm, a corrected absorbance signal specific to mRNA detection was obtained. The validity of such an approximation was validated using the Mie scattering theory (Fig. S10).

### Small Angle X-ray Scattering

The CoSAXS beamline at MAX IV has been constructed to deliver a high X-ray flux, 1013 photons/s at 12.4 keV, at a wavelength of *λ* = 0.99 Å. The data was collected on an Eiger2 4 M (Dectris) detector positioned at 3.5 m from the sample position and within an evacuated flight tube. The *q*-range accessible was 3.5 × 10−2 < *q* < 2.5 nm−1 where *q* = 4πsin(θ)/*λ*, where 2θ corresponds to the scattering angle and *λ* the X-ray wavelength. The flow cell in the Biocube of the CoSAXS beamline consists of a 0.98 mm inner-diameter quartz capillary, with a 10 µm wall thickness.

### AF4-SAXS fractogram data handling

The AF4-SAXS fractograms were collected at an acquisition rate of 1 frame/s, i.e. integration time 1 s. From the on-line SAXS acquisition, data in the fractograms was divided into 13 evenly distributed 30 s-fractions over the elution peaks, using CHROMIXS software from the ATSAS suite.^57,58^ Buffer frames were selected from a region of the elution where only carrier liquid elutes and their averaged signal was subtracted to the data.

### Indirect Fourier transformation method and Ab initio reconstruction

To carry out *p(r)* analysis, the GNOM software was employed within the BioXTAS RAW 2.3.0 suite.^41,59^ Guinier fits were performed according to the following specifications: *q*_min_\**R*_g_ < 1, *q*_max_\**R*_g_ < 1.3, and residuals demonstrating a relatively flat distribution around zero. *D*_max_, which represents the maximum linear dimension of the particle, is achieved by observing the characteristic decline of the *p(r)* curve to zero. To ensure the convergence of the model, the *p(r)* function was not force to zero at *D*_max_ and the regularization constant was modified to yield a realistic fit while minimizing the error. Once the *p(r)* was calculated for each scattering curve, low-resolution reconstruction of the LNP morphology was performed using the DAMMIN software.^60^ For each data file, 15 models were reconstructed using the “Slow” mode, P1 symmetry, and no pre-defined anisometry. Models were filtered and averaged to obtain the final reconstruction.

### Model-based analysis

The analysis of SAXS data employing mathematical models was performed using SasView 5.0.3.^61^ The form factor of the scatterers was modelled using a core-shell ellipsoid model to describe the overall particle structure,^45^ and a Teubner-Strey model to account for its internal organization.^39^ Sample polydispersity was taken into account by smearing the model using a Schulz function.^62^ Only polydispersity in the equatorial radius of the particle was found to improve the quality of the fit and the optimal polydispersity values ranged between 0.12 and 0.16. Data were fitted employing the Levenberg-Marquardt algorithm with a tolerance of 1.5×10^-8^ and uncertainties associated to the fitted parameters are derived from the mathematical analysis of the data.

### Electron Density Reconstruction Mapping Using DENSS

Following the *p(r)* calculations, files were submitted to DENSS web server for reconstruction. Twenty reconstructions of electron density mapping were performed in membrane mode, with *D*_max_ defined from the *p(r)* analysis and using the “Membrane” mode including negative-contrast components. Reconstruction alignment was achieved via the web server. Maps were visualized using PyMOL 3.1.6.1 Molecular Graphics System (Schrödinger, LLC. New York, NY). A volume object was generated, and six electron density contour levels were employed. The sigma (σ) level represents the standard deviation units above the average electron density value for the individual map. The absolute electron density corresponding to sigma levels can vary between maps; therefore, variance in electron density should only be interpreted within a single map, rather than compared across maps.

### Cryo-EM

All data was acquired using a Titan Krios G2 microscope (ThermoFisher Scientific) operating at 300 keV equipped with a Falcon 4i direct electron detector and a Selectris-X energy filter (ThermoFisher Scientific). 3 μL of N/P=3 or N/P=6 LNPs were applied to glow-discharged Quantifoil® 1.2/1.3 + 2 nm continuous carbon grids. The grids were blotted for 3 s at 4 °C and 100% humidity and plunge frozen into liquid ethane using a Vitrobot Mark IV (Thermo Fisher Scientific). All micrographs were collected using EPU software (ThermoFisher Scientific) at a nominal magnification of 81,000x (calibrated pixel size of 1.514 Å, FoV of 620 × 620 nm), a cumulative electron exposure of 50 e/Å^2 and a defocus range of -1 to -3.5. Micrographs were subsequently subjected to motion correction (in 5×5 patches) and CTF estimation using CryoSPARC 4.7.0 (Structura).

### Micrograph analysis

Representative images were randomly selected for qualitative analysis, and a total of 469 particles (N/P=3) and 507 particles (N/P=6) were analyzed. Particle morphology was evaluated using Fiji (ImageJ) following contrast normalization.^43^ Particles were categorized as either spherical, spherical bleb, or anisotropic bleb, with total counts per category recorded. Using the freehand selections tool, the outer particle boundaries for spherical blebs and anisotropic blebs were manually outlined, and the total particle areas recorded. For the same particles, individual blebs were separately outlined and quantified as individual bleb areas. For particles displaying multiple blebs, the total bleb area was calculated as the sum of its individual bleb areas.

### Nanoflow cytometry

Single-particle measurements of LNPs were performed using the Nanoanalyzer (NanoFCM Co.), which contain 488 and 647 nm lasers, each with bandpass filters of 525/40 and 670/30 for fluorescence detection. A quality control (QC) 250 nm silica bead solution of a fixed particle concentration (QC Beads, NanoFCM Co., Ltd) and a silica bead solution with sizes ranging from 68-155 nm (68-155 nm, NanoFCM Co., Ltd) were used to calibrate the instrument for nanoparticle concentration and size, respectively. When analyzing the QC beads, the detectors are aligned to result in side scattering events between 1.5-2k, FITC channel events between 1.5-2k and PC5 channel events between 500-1500 (20 mW laser power, both lasers), as per the company instruction manual. Both calibration solutions, LNPs, cleaning solution, and SYTO-9 dye were diluted in 20 mM Tris-EDTA (TE) buffer, which was filtered with a 0.1 µm filter prior to use. Fresh HPLC-grade MilliQ water (Fischer Scientific) was used as sheath fluid for each experiment. The detection threshold was based on automatic threshold assessment in the NanoFCM software.

The membrane-permeable RNA-binding SYTO-9 dye (ThermoFisher) was diluted 150-fold and kept in the dark. The LNPs were incubated with the diluted SYTO-9 dye in a LNP:SYTO-9 9:1 vol:vol ratio for 15 minutes in the dark. Following incubation, the mixture was diluted to exhibit between 2000-12000 events per minute. A blank measurement of TE-buffer was subtracted to all LNP samples. To assess the percentage of LNP-loading, a scatter plot of FITC-A/SS-S was made, which showed fluoresence-positive (loaded-LNPs) and fluoresence-negative (empty LNPs). To assess the mean RNA copy number, FITC-triggering was utilized to illuminate the fluorescence intensity of free RNA and loaded-LNPs, both stained with SYTO-9. The mRNA copy number was therefore calculated by dividing the fluorescent intensities of each LNP in the loaded-LNP population with the mean fluorescence intensity of free RNA. The RNA copy number distribution in LNPs was normalized as percentage of total events and plotted as a histogram.

### In-vitro transfection assays

HEK293 cells were obtained from ATCC (Cat# CRL-1573). Cells were cultured in DMEM (Dulbecco’s Modified Eagle’s Medium, high glucose, GlutaMAX supplement, pyruvate; Thermo Fisher Scientific/Gibco, Cat# 31966021) supplemented with 10% heat-inactivated fetal bovine serum (FBS; Thermo Fisher Scientific/Gibco, Cat# A5670801) and 1% penicillin-streptomycin (Thermo Fisher Scientific/Gibco, Cat# 15140122). Cultures were maintained at 37 °C in a humidified atmosphere with 5% CO₂. For gentle detachment during passaging, TrypLE Express Enzyme (Thermo Fisher Scientific, Cat# 12604013) was used.

Cultured HEK293 cells were detached and counted using a Countess 3 Automated Cell Counter (Thermo Fisher Scientific) by mixing 10 µL of the cell suspension with 10 µL Trypan Blue Stain (Thermo Fisher Scientific, Cat# T10282). Cell viability was assessed (100%), and the suspension was diluted to 25,000 cells per well in 100 µL and seeded into a 96-well PhenoPlate (Revvity, Cat# 6055300). Cells were incubated for 24 h at 37 °C and 5% CO₂. Following incubation, the medium was replaced with 100 µL of fresh culture medium containing 0.5 µg/mL Hoechst 33342 (Thermo Fisher Scientific, Cat# 62249) and mRNA-LNPs at doses of 25, 50, 100, or 200 ng/well. The mRNA-LNPs were diluted in PBS (Thermo Fisher Scientific, Cat# 10010015) and culture medium containing Hoechst 33342 to achieve a final FBS concentration of 9.75%. Controls and Imaging: PBS was used as a negative control. As a positive control, 50 ng/well EGFP mRNA (Trilink, Cat# L-7601-1000) was transfected using Lipofectamine MessengerMax (Thermo Fisher Scientific, Cat# LMRNA008). Brightfield and fluorescence imaging were performed using an Operetta CLS confocal microscope (Revvity).

Statistical differences in transfection efficiency between formulations were validated using a two-way analysis of variance (ANOVA) with formulation (N/P 3 and N/P 6) and dose (25, 50, 100, and 200 ng mRNA per well; or 0.58, 1.17, and 2.33 µg total lipid per well) as fixed factors. Prior to analysis, data from three independent replicates per condition were verified to satisfy ANOVA assumptions of homoscedasticity and approximate normality. To assess pairwise diffe∫rences between formulations at each dose, Welch’s *t*-tests were performed. Resulting *p*-values were corrected for multiple comparisons using the Holm method. Statistical significance was defined as *p* < 0.05 after correction. All analyses were carried out in Python using the SciPy and statsmodels packages.

## Acknowledgements

We acknowledge funding from the Swedish Research Council (VR) grant no. 2021-04667_VR. Funding for AF4 equipment was partially obtained from the Royal Physiographic Society in Lund, Sweden and by strategic funding from the Faculty of Engineering, Lund University. The conducted research was partially supported by The University of Texas at Austin and the National Institute of General Medical Sciences at the National Institutes of Health under award R35 GM154984. A.S.-F. is thankful for the financial support of Spanish grants RYC2022-037909-I and PID2022-141673OA-I00 funded by MCIN/AEI and by “ERDF A way of making Europe”. We acknowledge MAX IV Laboratory for time on Beamline CoSAXS under Proposal #: 20250974. Research conducted at MAX IV, a Swedish national user facility, is supported by the Swedish Research council under contract 2018-07152, the Swedish Governmental Agency for Innovation Systems under contract 2018-04969, and Formas under contract 2019-02496. We thank Berit Bergerud Dilling and Sanne Brandt Nielsen, Principal Laboratory Technicians at Novo Nordisk, for the expert production of LNPs and characterization by DLS and Ribogreen and the expert execution of in vitro assays and subsequent analysis, respectively. We also thank Prof. Thomas Grant at the University of Buffalo for helpful discussions on DENSS.

## Data availability

All data supporting the findings of this study is presented in the manuscript and Supplementary Information file. Source data is available from the corresponding author upon reasonable request.

## References

1 Opalinska, J. B. & Gewirtz, A. M. Nucleic-acid therapeutics: basic principles and recent applications. Nat Rev Drug Discov 1, 503–514 (2002).

2 Lostalé-Seijo, I. & Montenegro, J. Synthetic materials at the forefront of gene delivery. Nature Reviews Chemistry 2, 258–277 (2018).

3 Kulkarni, J. A. et al. The current landscape of nucleic acid therapeutics. Nat. Nanotechnol. 16, 630–643 (2021).

4 Hou, X., Zaks, T., Langer, R. & Dong, Y. Lipid nanoparticles for mRNA delivery. Nat Rev Mater 6, 1078–1094 (2021).

5 Yap, S. L. et al. The Internal Nanostructure of Lipid Nanoparticles Influences Their Diverse Cellular Uptake Pathways. Small 21, e2500903 (2025).

6 Mulligan, M. J. et al. Phase I/II study of COVID-19 RNA vaccine BNT162b1 in adults. Nature 586, 589–593 (2020).

7 Caselli, L., Conti, L., De Santis, I. & Berti, D. Small-angle X-ray and neutron scattering applied to lipid-based nanoparticles: Recent advancements across different length scales. Adv. Colloid Interface Sci. 327, 103156 (2024).

8 Udepurkar, A. et al. Structure and Morphology of Lipid Nanoparticles for Nucleic Acid Drug Delivery: A Review. ACS Nano 19, 21206–21242 (2025).

9 Eygeris, Y., Gupta, M., Kim, J. & Sahay, G. Chemistry of Lipid Nanoparticles for RNA Delivery. Acc. Chem. Res. 55, 2–12 (2022).

10 Mehraji, S. & DeVoe, D. L. Microfluidic synthesis of lipid-based nanoparticles for drug delivery: recent advances and opportunities. Lab Chip 24, 1154–1174 (2024).

11 Tenchov, R., Bird, R., Curtze, A. E. & Zhou, Q. Lipid Nanoparticles horizontal line From Liposomes to mRNA Vaccine Delivery, a Landscape of Research Diversity and Advancement. ACS Nano 15, 16982–17015 (2021).

12 Cárdenas, M., Campbell, R. A., Yanez Arteta, M., Lawrence, M. J. & Sebastiani, F. Review of structural design guiding the development of lipid nanoparticles for nucleic acid delivery. Current Opinion in Colloid & Interface Science 66, 101705 (2023).

13 Thelen, J. L. et al. Morphological Characterization of Self-Amplifying mRNA Lipid Nanoparticles. ACS Nano 18, 1464–1476 (2024).

14 Sabnis, S. et al. A Novel Amino Lipid Series for mRNA Delivery: Improved Endosomal Escape and Sustained Pharmacology and Safety in Non-human Primates. Mol. Ther. 26, 1509–1519 (2018).

15 Li, Z., et al. Acidification-Induced Structure Evolution of Lipid Nanoparticles Correlates with Their In Vitro Gene Transfections. ACS Nano 17, 979–990 (2023).

16 Devos, C. et al. Manufacturing mRNA-Loaded Lipid Nanoparticles with Precise Size and Morphology Control. ACS Nano 19, 33991–34002 (2025).

17 Gilbert, J. et al. Evolution of the structure of lipid nanoparticles for nucleic acid delivery: From in situ studies of formulation to colloidal stability. J. Colloid Interface Sci. 660, 66–76 (2024).

18 Simonsen, J. B. A perspective on bleb and empty LNP structures. J Control Release 373, 952–961 (2024).

19 Borjesdotter, A. M. et al. Lipid nanoparticle properties explored using online asymmetric flow field-flow fractionation coupled with small angle X-ray scattering: Beyond average characterisation. Int J Pharm 668, 124940 (2025).

20 Jia, X. et al. Enabling online determination of the size-dependent RNA content of lipid nanoparticle-based RNA formulations. J Chromatogr B Analyt Technol Biomed Life Sci 1186, 123015 (2021).

21 Patel, S. et al. Naturally-occurring cholesterol analogues in lipid nanoparticles induce polymorphic shape and enhance intracellular delivery of mRNA. Nat. Commun. 11, 983 (2020).

22 Philipp, J. et al. pH-dependent structural transitions in cationic ionizable lipid mesophases are critical for lipid nanoparticle function. Proc. Natl. Acad. Sci. U. S. A. 120, e2310491120 (2023).

23 Pattipeiluhu, R. et al. Liquid crystalline inverted lipid phases encapsulating siRNA enhance lipid nanoparticle mediated transfection. Nat. Commun. 15, 1303 (2024).

24 Dao, H. M. et al. Characterization of mRNA Lipid Nanoparticles by Electron Density Mapping Reconstruction: X-ray Scattering with Density from Solution Scattering (DENSS) Algorithm. Pharm. Res. 41, 501–512 (2024).

25 Padilla, M. S. et al. Elucidating lipid nanoparticle properties and structure through biophysical analyses. Nat. Biotechnol. (2025).

26 Eygeris, Y., Patel, S., Jozic, A. & Sahay, G. Deconvoluting Lipid Nanoparticle Structure for Messenger RNA Delivery. Nano Lett. 20, 4543–4549 (2020).

27 Haque, M. A., Shrestha, A., Mikelis, C. M. & Mattheolabakis, G. Comprehensive analysis of lipid nanoparticle formulation and preparation for RNA delivery. Int J Pharm X 8, 100283 (2024).

28 Li, S. et al. Payload distribution and capacity of mRNA lipid nanoparticles. Nat. Commun. 13, 5561 (2022).

29 Hammel, M. et al. Correlating the Structure and Gene Silencing Activity of Oligonucleotide-Loaded Lipid Nanoparticles Using Small-Angle X-ray Scattering. ACS Nano 17, 11454–11465 (2023).

30 Garaizar, A. et al. Toward understanding lipid reorganization in RNA lipid nanoparticles in acidic environments. Proc. Natl. Acad. Sci. U. S. A. 121, e2404555121 (2024).

31 Bolinsson, H., Soderberg, C., Herranz-Trillo, F., Wahlgren, M. & Nilsson, L. Realizing the AF4-UV-SAXS on-line coupling on protein and antibodies using high flux synchrotron radiation at the CoSAXS beamline, MAX IV. Anal. Bioanal. Chem. 415, 6237–6246 (2023).

32 Da Vela, S. et al. AF4-to-SAXS: expanded characterization of nanoparticles and proteins at the P12 BioSAXS beamline. J. Synchrotron Radiat. 32, 971–985 (2025).

33 Wahlund, K. G. & Giddings, J. C. Properties of an asymmetrical flow field-flow fractionation channel having one permeable wall. Anal. Chem. 59, 1332–1339 (1987).

34 Wahlund, K.-G. & Nilsson, L. in Field-Flow Fractionation in Biopolymer Analysis (eds S. Kim R. Williams & Karin D. Caldwell) Ch. Chapter 1, 1–21 (Springer Vienna, 2012).

35 Graewert, M. A. et al. Quantitative size-resolved characterization of mRNA nanoparticles by in-line coupling of asymmetrical-flow field-flow fractionation with small angle X-ray scattering. Sci. Rep. 13, 15764 (2023).

36 Moon, M. H., Kwon, H. & Park, I. Stopless flow injection in asymmetrical flow field-flow fractionation using a frit inlet. Anal. Chem. 69, 1436–1440 (1997).

37 Fuentes, C. et al. Comparison between conventional and frit-inlet channels in separation of biopolymers by asymmetric flow field-flow fractionation. Analyst 144, 4559–4568 (2019).

38 Bolinsson, H. et al. Sum-weighted casein micelle AF4-UV-SAXS data disentangled - A new method for characterization and evaluation of widely size distributed samples. Food Hydrocolloids 166, 111377 (2025).

39 Teubner, M. & Strey, R. Origin of the scattering peak in microemulsions. J. Chem. Phys. 87, 3195–3200 (1987).

40 Glatter, O. Determination of particle-size distribution functions from small-angle scattering data by means of the indirect transformation method. J. Appl. Crystallogr. 13, 7–11 (1980).

41 Hopkins, J. B. BioXTAS RAW 2: new developments for a free open-source program for small-angle scattering data reduction and analysis. J. Appl. Crystallogr. 57, 194–208 (2024).

42 Mertens, H. D. & Svergun, D. I. Structural characterization of proteins and complexes using small-angle X-ray solution scattering. J Struct Biol 172, 128–141 (2010).

43 Schindelin, J., et al. Fiji: an open-source platform for biological-image analysis. Nat. Methods 9, 676–682 (2012).

44 Li, H. et al. Mesoscopic Structure of Lipid Nanoparticles Studied by Small-Angle X-Ray Scattering: A Spherical Core-Triple Shell Model Analysis. Membranes (Basel) 15, 153 (2025).

45 Berr, S. S. Solvent isotope effects on alkytrimethylammonium bromide micelles as a function of alkyl chain length. J. Phys. Chem. 91, 4760–4765 (2002).

46 Kulkarni, J. A. et al. On the Formation and Morphology of Lipid Nanoparticles Containing Ionizable Cationic Lipids and siRNA. ACS Nano 12, 4787–4795 (2018).

47 Leung, A. K., Tam, Y. Y., Chen, S., Hafez, I. M. & Cullis, P. R. Microfluidic Mixing: A General Method for Encapsulating Macromolecules in Lipid Nanoparticle Systems. J. Phys. Chem. B 119, 8698–8706 (2015).

48 Louro, A. F. et al. Engineering Hybrid Extracellular Vesicles for Functional mRNA Delivery. Adv. Funct. Mater. n/a, e09636 (2025).

49 Chen, X. et al. Structural characterization of mRNA lipid nanoparticles (LNPs) in the presence of mRNA-free LNPs. J Control Release 386, 114082 (2025).

50 Grant, T. D. Ab initio electron density determination directly from solution scattering data. Nat. Methods 15, 191–193 (2018).

51 Wang, R. et al. Understanding the self-assembly and molecular structure of mRNA lipid nanoparticles at real size: Insights from the ultra-large-scale simulation. Int J Pharm 670, 125114 (2025).

52 Liao, S. et al. Transfection Potency of Lipid Nanoparticles Containing mRNA Depends on Relative Loading Levels. ACS Appl. Mater. Interfaces 17, 3097–3105 (2025).

53 Cabry, C. P. et al. Exploring the bulk-phase structure of ionic liquid mixtures using small-angle neutron scattering. Faraday Discuss. 206, 265–289 (2018).

54 Wilhelmy, C. et al. Direct structural investigation of pH responsiveness in mRNA lipid nanoparticles: Refining paradigms. J Control Release 384, 113848 (2025).

55 Ventouri, I. K., Loeber, S., Somsen, G. W., Schoenmakers, P. J. & Astefanei, A. Field-flow fractionation for molecular-interaction studies of labile and complex systems: A critical review. Anal. Chim. Acta 1193, 339396 (2022).

56 Doucet, M. et al. SasView version 6.1.0 (v6.1.0), 2025).

57 Manalastas-Cantos, K. et al. ATSAS 3.0: expanded functionality and new tools for small-angle scattering data analysis. J. Appl. Crystallogr. 54, 343–355 (2021).

58 Panjkovich, A. & Svergun, D. I. CHROMIXS: automatic and interactive analysis of chromatography-coupled small-angle X-ray scattering data. Bioinformatics 34, 1944–1946 (2018).

59 Svergun, D. I. Determination of the regularization parameter in indirect-transform methods using perceptual criteria. J. Appl. Crystallogr. 25, 495–503 (1992).

60 Svergun, D. I. Restoring low resolution structure of biological macromolecules from solution scattering using simulated annealing. Biophys. J. 76, 2879–2886 (1999).

61 Doucet, M., et al. SasView version 5.0.3, <https://zenodo.org/record/3930098> (2020).

62 Kotlarchyk, M. & Chen, S.-H. Analysis of small angle neutron scattering spectra from polydisperse interacting colloids. J. Chem. Phys. 79, 2461–2469 (1983).

