## Supplementary Information for "Structural heterogeneity in mRNA-LNP subpopulations revealed by AF4-SAXS: implications for cargo loading and cell transfection"

[a]Centro Singular de Investigación en Química Biolóxica e Materiais Moleculares (CIQUS), Departamento de Enxeñaría Química, Universidade de Santiago de Compostela, Santiago de Compostela, Spain.
[b]Department of Biomedical Engineering, The University of Texas at Austin, Austin, Texas 78712, United States
[c]Department of Process and Life Science Engineering, Lund University, Lund, Sweden
[d]Nucleic Acid Research, Novo Nordisk A/S, Måløv, Denmark.
[e]Research Centres of Excellence, Novo Nordisk A/S, Måløv, Denmark.
[f]Walker Department of Mechanical Engineering, The University of Texas at Austin, Austin, Texas 78712, United States
[g]McKetta Department of Chemical Engineering, The University of Texas at Austin, Austin, Texas 78712, United States
[h]MAX IV Laboratory, Lund University, Lund, Sweden
[i]Drug Product Research, Novo Nordisk A/S, Måløv, Denmark.
[j]Nucleic Acid Research, Novo Nordisk, Lexington, Massachusetts, United States.
*These authors contributed equally: Adrian Sanchez-Fernandez, Keira A. Donnelly.

### 1) Results from the ensemble-averaged SAXS analysis of LNPs

Table S1 Results from the model-based analysis of SAXS data from the unfractionated LNPs formulated at N/P = 3. The model used to fit the data was a combination of the core-shell ellipsoid and the Teubner-Strey model. The parameters extracted from the modelling are: concentration of LNPs (*[LNP]*), equatorial radius of the particle ($r_{eq}$), thickness of the shell ($t_{shell}$), aspect ratio (AR), scattering length densities of the core ($SLD_{core}$) and shell ($SLD_{shell}$), periodicity ($d$), correlation length ($\xi$), and polydispersity in the equatorial dimension (*PD*).

| [LNP] / $\times 10^{15}$ m$^{-3}$ | $r_{eq}$ / nm | $t_{shell}$ / nm | AR | $SLD_{core}$ / $\times 10^{-6}$ Å | $SLD_{shell}$ / $\times 10^{-6}$ Å | d / nm | $\xi$ / nm | PD |
|---|---|---|---|---|---|---|---|---|
| 54.7±0.3 | 30.9±0.4 | 11.9±0.2 | 2.07±0.12 | 12.5±0.2 | 8.56±0.08 | 5.3±0.2 | 2.9±0.3 | 0.15 |

### 2) Structural heterogeneity observed by cryo-EM

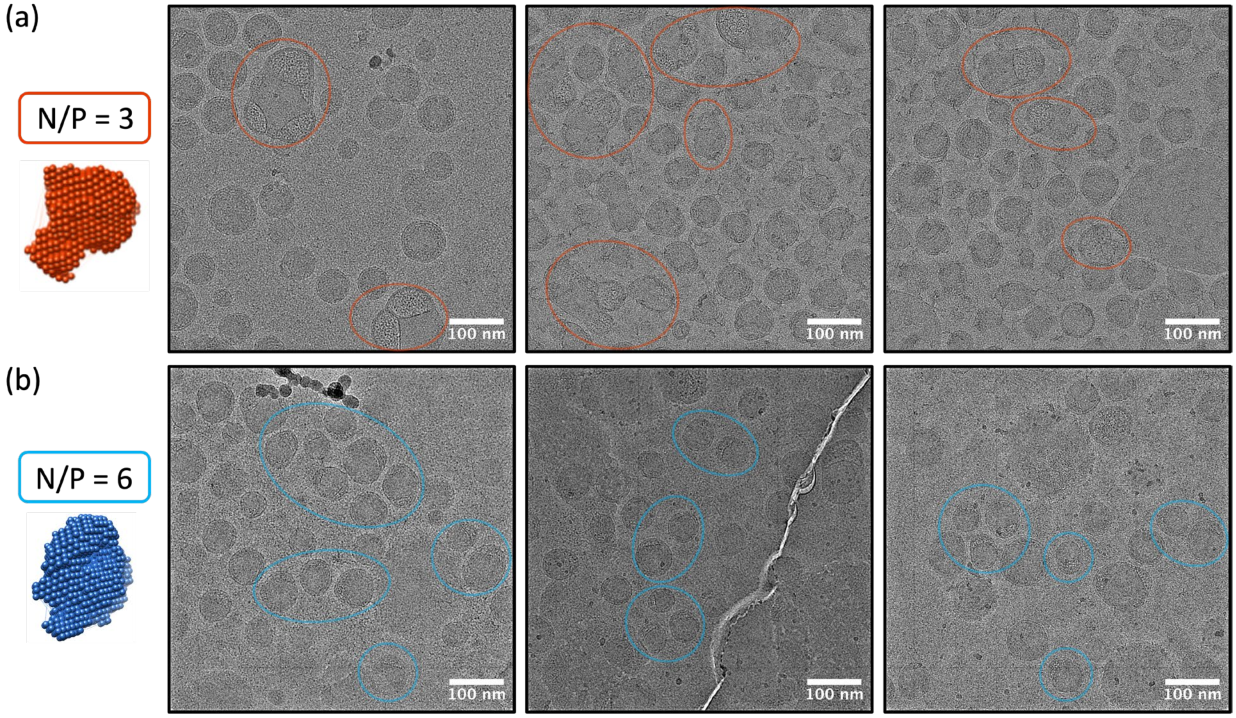

**Fig. S1** Representative cryo-EM micrographs show differences in bleb morphologies for N/P = 3 (larger, more anisotropic) and N/P = 6 (smaller, less anisotropic) particles. Circles highlight particles corresponding to the specified morphology.

#### 3) Spectroscopic signals from the fractions acquired by AF4

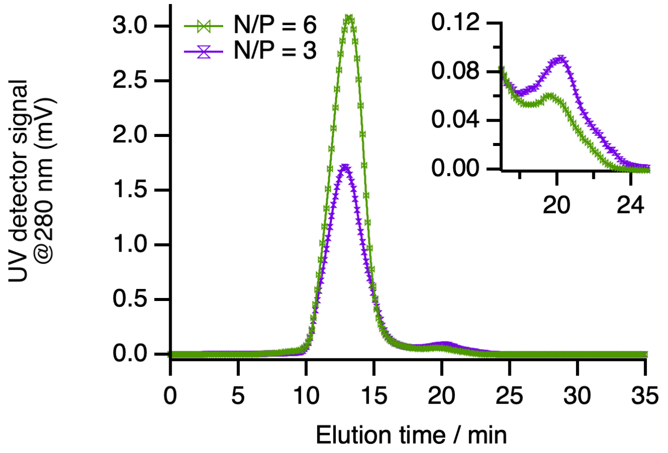

**Fig. S2** UV signal acquired at 280 during the AF4 fractionation of LNPs formulated at N/P = 3 and 6, as indicated in the legend of the graph. The inset shows the signal from the tail of the peak.

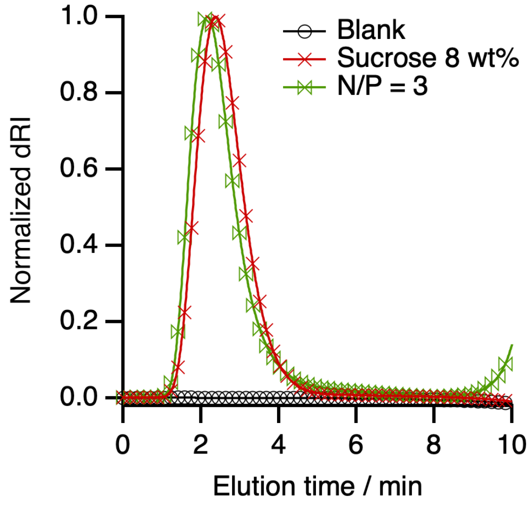

**Fig. S3** Fractogram with the comparison of the dRI signal in the early peak (≈ 2.5 min) between an LNP sample, the buffer, and the carrier liquid, as indicated in the legend of the graph.

Considering the high dRI signal, the increase in the scattered intensity, and the absence of UV signal (**Fig. 1**), this peak was attributed to the elution of formulation excipients, as confirmed by injection of a sucrose solution (8 wt%) into the AF4 channel. The injection/elution of a blank is also shown, ruling out systematic instrument contribution.

### 4) Elution times for the AF4 fractions

**Table S2** Elution time windows for AF4 fractions used for SAXS binning.

| N/P = 3 | | | N/P = 6 | | |
|---|---|---|---|---|---|
| **Fraction** | **Elution start / min** | **Elution end / min** | **Fraction** | **Elution start / min** | **Elution end / min** |
| **1** | 10.2 | 10.6 | **1** | 10.4 | 10.8 |
| **2** | 10.7 | 11.1 | **2** | 10.8 | 11.2 |
| **3** | 11.1 | 11.5 | **3** | 11.3 | 11.7 |
| **4** | 11.6 | 12.0 | **4** | 11.7 | 12.1 |
| **5** | 12.1 | 12.5 | **5** | 12.2 | 12.6 |
| **6** | 12.6 | 13.1 | **6** | 12.7 | 13.1 |
| **7** | 13.1 | 13.6 | **7** | 13.2 | 13.6 |
| **8** | 13.7 | 14.1 | **8** | 13.7 | 14.2 |
| **9** | 14.2 | 14.7 | **9** | 14.3 | 14.7 |
| **10** | 14.8 | 15.2 | **10** | 14.9 | 15.3 |
| **11** | 15.3 | 15.8 | **11** | 15.4 | 15.9 |
| **12** | 15.8 | 16.3 | **12** | 16.1 | 16.5 |
| **13** | 16.4 | 16.8 | **13** | 16.7 | 17.2 |

### 5) SAXS data analysis of the fractionated samples

**Table S3** Molecular volumes ($v_m$), scattering lengths ($\sum b_i$), and scattering length densities (SLDs) of the components present in the systems.

| Component | Molecular volume / Å3 | Scattering length / fm | SLD / x10-6 Å2 |
|---|---|---|---|
| **Buffer** | 30 | 28.2 | 9.40 |
| **DSPC** | 1235.2 | 1327 | 9.31 |
| **Cholesterol** | 609.1 | 630 | 9.67 |
| **Lipid 5** | 1020.8 | 1300 | 7.85 |
| **DMG-PEG2k** | 3878.7 | 3630 | 10.7 |
| **mRNA** | - | - | 15.5 |

The SLD of each component was calculated using the following equation:

$$SLD = \frac{\sum b_i}{v_m}$$

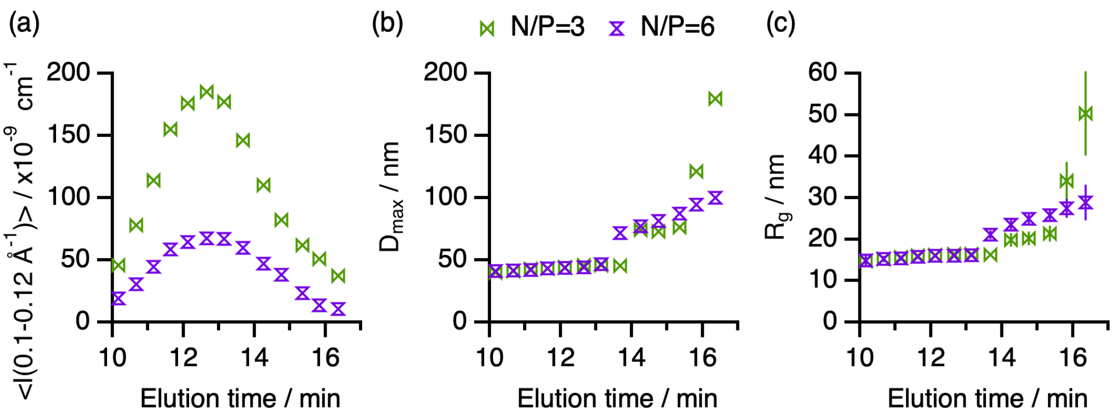

**Fig. S4** Results from SAXS-based analysis of particle structure across fractions for N/P = 3 and N/P = 6: (a) changes in the average peak intensity for N/P = 3 and N/P = 6, (b) changes in Dmax, and (c) changes in Rg. Error bars represent the standard deviation to the averages. Where not seen, error bars are within the markers.

The parameters extracted from the fits allowed us to determine the volume fraction of mRNA in the particle core using the following equation:

$$SLD_{core} = x_{lipids}SLD_{lipids} + x_{mRNA}SLD_{mRNA} + x_{buffer}SLD_{buffer}$$

where *x* corresponds to the volume fraction of each component in the particle core. The scattering length density of the lipids at the core ($SLD_{lipids}$) was calculated based on the assumption that 85% of those is Lipid 5 and 15% is cholesterol.[1] The volume fraction of buffer at the particle core was fixed to 0.32.[2] These compositions were assumed to remain unchanged for the different fractions. From this equation, the fraction of mRNA at the core ($x_{mRNA}$) was determined from the fitted $SLD_{core}$. Although our assumptions may introduce some uncertainties in the calculations, our results broadly agree with previous reports on the average volume fraction of mRNA at the particle core.[1]

### 6) Micrograph analysis of LNPs

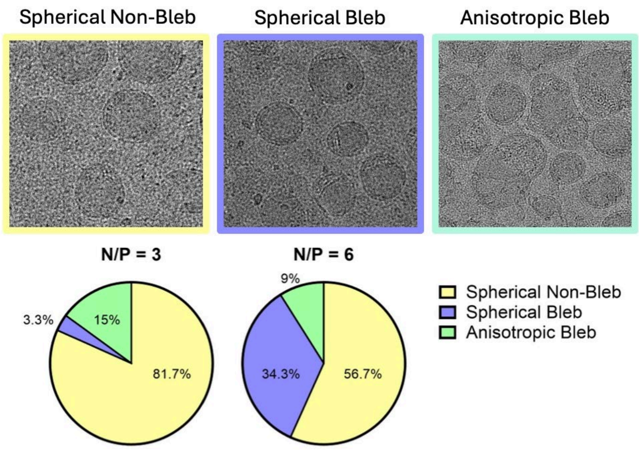

**Fig. S5** Cryo-EM validation of particle heterogeneity showing the presence of different structures in the formulations.

### 7) Single-particle analysis by nanoflow cytometry

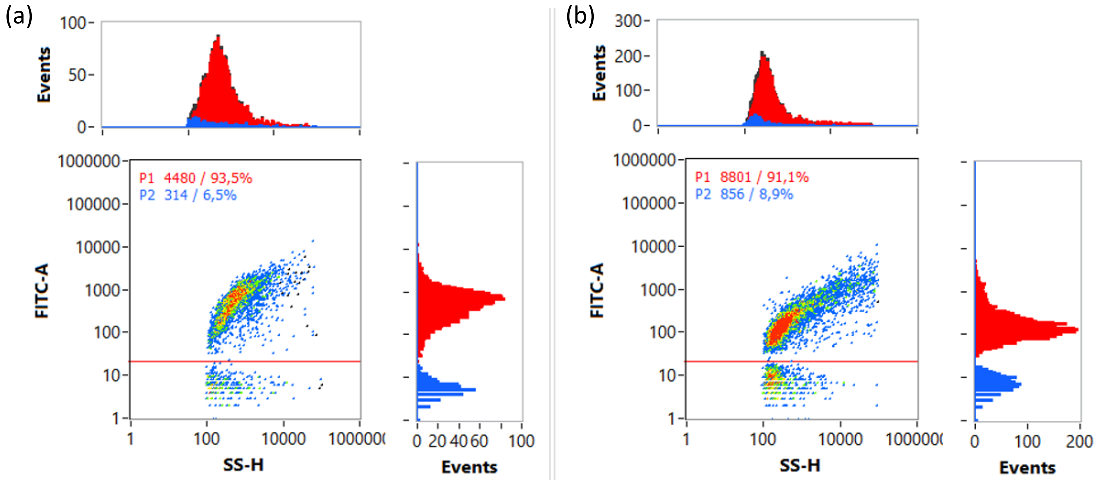

**Fig. S6** Representative nanoflow cytometry plots showing the fluorescent signal of SYTO-9 labelling of mRNA (FITC-A channel) against the optical size (SS-H channel) of LNPs for individual events. The red line represents the gate vale used to differentiate between fluorescent-negative (below) and fluorescence-positive (above) LNPs. LNP concentration was adjusted to acquire between 2000 and 12000 events per sample.

Side scattering and fluorescence of RNA-LNPs was measured simultaneously using nanoflow cytometry by incubating the RNA-LNPs with a membrane permeable dye, SYTO-9. The FITC triggering channel was used to gate out fluorescent negative events (empty LNPs), allowing to determine fluorescence-negative (empty) and fluorescence-positive (loaded) LNPs.

### 8) Peak deconvolution analysis of SAXS data

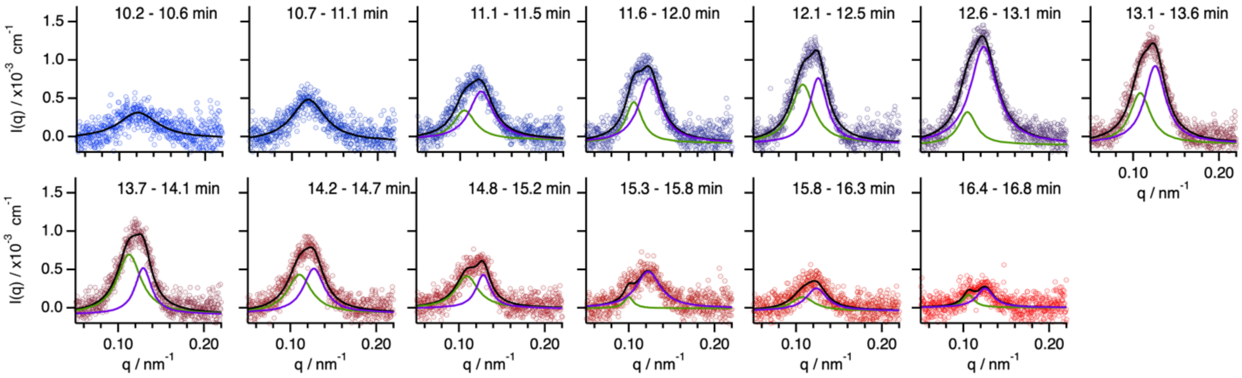

**Fig. S7** SAXS data (markers) and fits (lines) for the peak deconvolution approach. The plots correspond to the AF4 fractions with increasing elution time, as indicated in the legend of the graphs.

**Table S4** Fitting parameters obtained from peak deconvolution for N/P = 3 LNPs.

| Fraction | Peak 1 (low q) | | | Peak 2 (high q) | | |
|---|---|---|---|---|---|---|
| | Peak center (Å$^{-1}$) | Intensity (cm$^{-1}$) | d-spacing (nm) | Peak center (Å$^{-1}$) | Intensity (cm$^{-1}$) | d-spacing (nm) |
| 1 | - | - | - | 0.1214 | 0.0003 | 51.7390 |
| 2 | - | - | - | 0.1190 | 0.0006 | 52.8088 |
| 3 | 0.1048 | 0.0004 | 59.9426 | 0.1243 | 0.0006 | 50.5486 |
| 4 | 0.1058 | 0.0005 | 59.4042 | 0.1244 | 0.0008 | 50.5282 |
| 5 | 0.1077 | 0.0008 | 58.3668 | 0.1257 | 0.0009 | 49.9856 |
| 6 | 0.1046 | 0.0004 | 60.0457 | 0.1234 | 0.0013 | 50.9131 |
| 7 | 0.1082 | 0.0007 | 58.0916 | 0.1255 | 0.0010 | 50.0812 |
| 8 | 0.1126 | 0.0008 | 55.8158 | 0.1291 | 0.0006 | 48.6880 |
| 9 | 0.1104 | 0.0005 | 56.9232 | 0.1271 | 0.0006 | 49.4233 |
| 10 | 0.1091 | 0.0004 | 57.5700 | 0.1282 | 0.0004 | 49.0299 |
| 11 | 0.0998 | 0.0001 | 62.9515 | 0.1225 | 0.0005 | 51.2746 |
| 12 | 0.1076 | 0.0002 | 58.3776 | 0.1235 | 0.0003 | 50.8636 |
| 13 | 0.1059 | 0.0002 | 59.3201 | 0.1252 | 0.0003 | 50.1892 |

### 9) Reconstructions of the particle electron density map

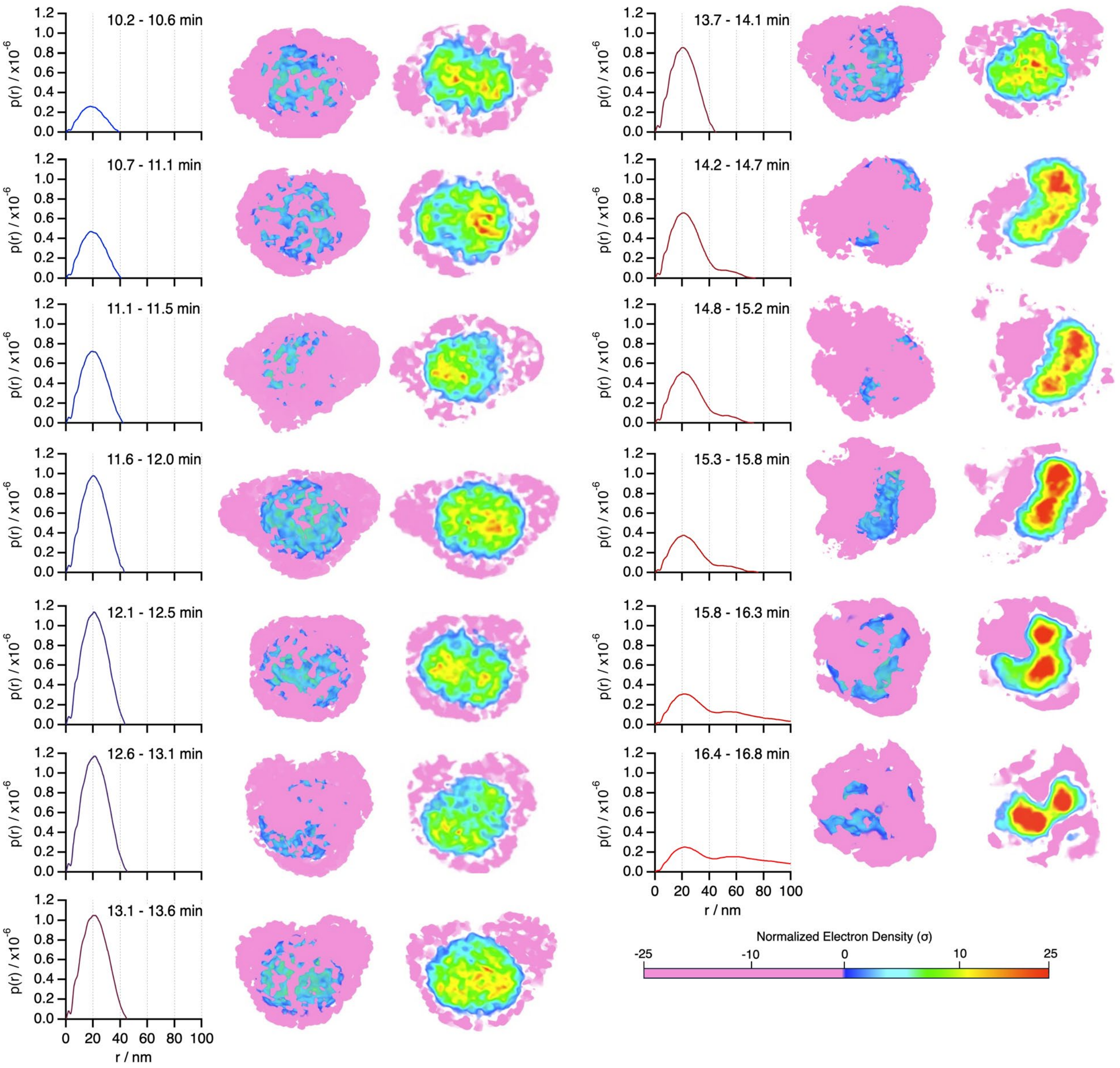

**Fig. S8** *p(r)* functions (left) and DENSS reconstructions showing the external (middle) and internal (right) views of electron density for N/P = 3 particles.

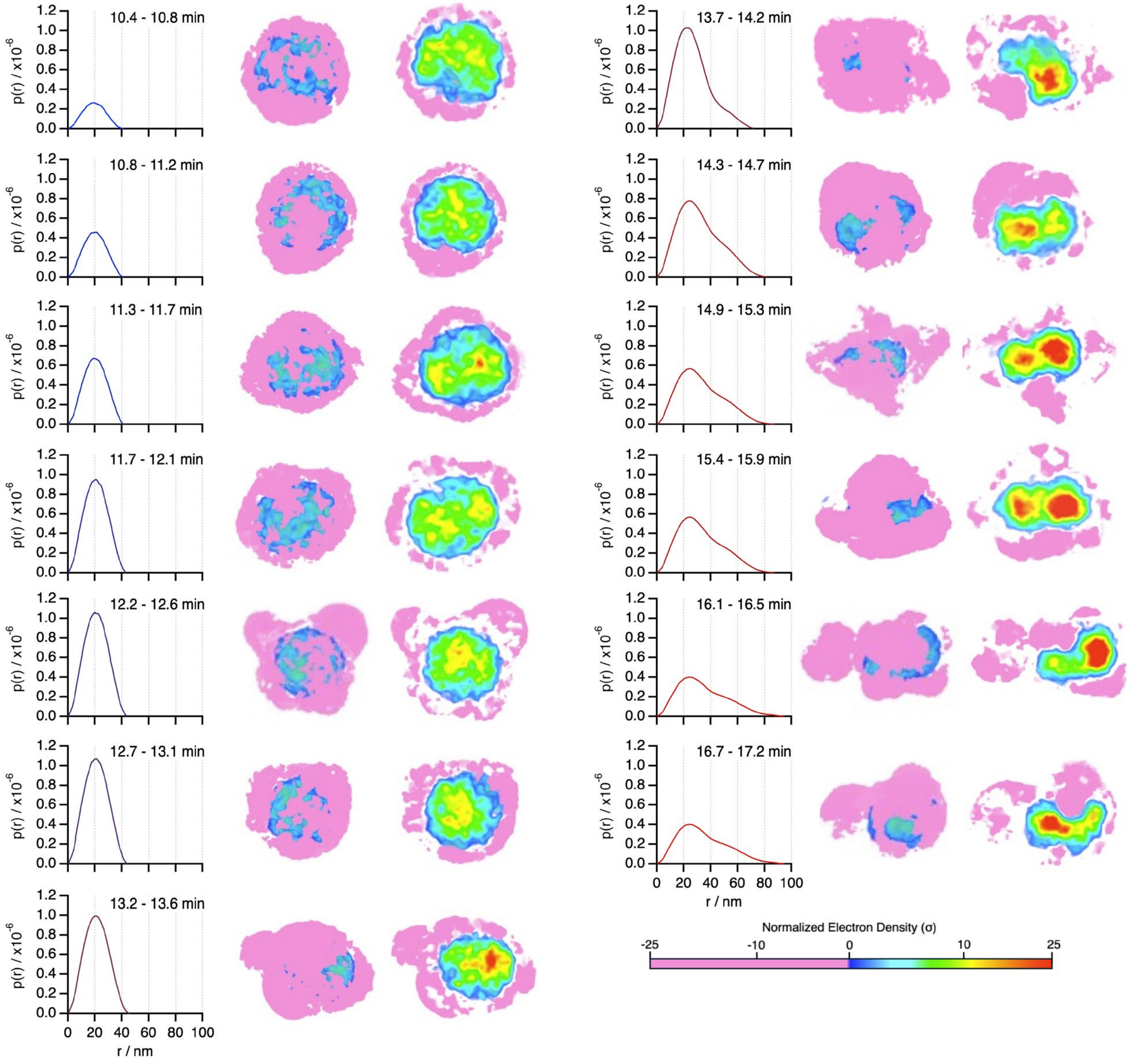

**Fig. S9** *p(r)* functions (left) and DENSS reconstructions showing the external (middle) and internal (right) views of electron density for N/P = 6 particles.

The results from the analysis show how the *p(r)* shift from unimodal to bimodal shapes, indicative of a transition from spherical morphology to anisotropic, bleb structures. This is further supported by the DENSS reconstructions. Internal views display the distribution of electron density within the particle and bleb compartmentalization, while external views highlight overall particle shape.

### 10) Mie theory scattering correction of the absorbance

Rayleigh scattering theory assumes that the particles' diameter (*d*) is much smaller, approx. 1/20, of the than the wavelength of the light ($\lambda$). By utilizing the Rayleigh-Gans-Debye (RGD) approximation, $|(n_{particles}/n_{medium})-1| << 1$ i.e. low contrast between the scatterer and the surrounding medium, Rayleigh theory can be extended to $d \leq \lambda$. For our system it is reasonable to assume that the RGD criterion is fulfilled.

By assuming a homogeneous sphere, the largest $R_g$ in Fig. 1 can be converted to a geometric diameter ($d$) through

$$d = \frac{2 \cdot R_g}{\sqrt{\frac{3}{5}}}$$

which gives us $d \approx 210$ nm, i.e. < 280 nm, suggesting that the approach is adequate.

The Rayleigh scattering approach was validated by determining the $A_{280}^{corr}$ for each AF4 fraction with Mie scattering theory using PyMieLab v1.0.[3] The Mie scattering efficiency (Qscat) was calculated and plotted in Figure S8(b) as a function of particle radius at two wavelengths: 280 nm and 340 nm for lipid nanoparticles of 1.45 refractive index.[4] A homogenous sphere was assumed.

Then, the ratio of Qscat(280)/ Qscat(340) was calculated from Mie theory to yield:

$$A_{280}^{corr} = A_{280}^{meas} - \frac{Q_{sca}(280)}{Q_{sca}(340)} A_{340}^{meas}$$

where the second term represents scattering only at 280 nm given the absorption at 340 nm is 0. The $A_{280}$ for each fraction was normalized for the total $A_{280}$ for all fractions as shown in Figure S8(b). The differences remained negligible even though the particle size reached approx. 70% the wavelength.

**Fig. S10** (a) Mie scattering efficiency ($Q_{scat}$) vs. particle radius at 280 nm and 340 nm. (b) Mie and Rayliegh scattering-corrected relative absorbances at 280 nm for N/P = 3.

### 11) References

1 Sebastiani, F. *et al.* Apolipoprotein E Binding Drives Structural and Compositional Rearrangement of mRNA-Containing Lipid Nanoparticles. *ACS Nano* **15**, 6709-6722 (2021).

2 Liu, H. *et al.* Mapping Hydration and Nanoarchitecture in mRNA-Loaded Lipid Nanoparticles Through Small-Angle Neutron Scattering. *Small Structures* **n/a**, e202500636 (2025).

3 Ma, D., Tuersun, P., Cheng, L., Zheng, Y. & Abulaiti, R. PyMieLab_V1.0: A software for calculating the light scattering and absorption of spherical particles. *Heliyon* **8**, e11469 (2022).

4 Meredith, S. A. *et al.* Model Lipid Membranes Assembled from Natural Plant Thylakoids into 2D Microarray Patterns as a Platform to Assess the Organization and Photophysics of Light-Harvesting Proteins. *Small* **17**, 2006608 (2021).
